# Basal Ganglia Dopamine Availability and Immune Activation Interact and Relate to Anhedonia Severity among Youth with Depression

**DOI:** 10.64898/2025.12.23.696320

**Authors:** Iris Ka-Yi Chat, James W. Murrough, Scott Russo, Finnegan Calabro, Beatriz Luna, Kalyan Tripathy, Tina Gupta, Zachary Brodnick, Megan M. Julien, Neil P. Jones, Neal D. Ryan, Helmet T. Karim, Rebecca B. Price, Jihui Diaz, Erika E. Forbes

## Abstract

Anhedonia emerges in adolescence, has putative substrates in neural reward and dopamine systems, and is a hallmark of poor course in depression. Psychoneuroimmunology models suggest altered immune and reward systems may jointly underlie its development. When dopamine availability (rather than function) is considered, immune activation may be understood as a moderator amplifying how lower dopamine levels translate into motivational deficits. This cross-sectional study analyzed data from 55 youth with current depression (M_age_=21.4 years; Female=75%). Plasma was assayed for relative quantification of 92 immune proteins, followed by dimensionality reduction via principal component (PC) analysis, forming five PC scores. Dopamine availability was indexed by basal ganglia tissue iron, quantified by MRI R2’, a measure detectable in youth. Anhedonia was assessed using the Snaith–Hamilton Pleasure Scale (SHAPS), thought to capture overall anhedonia, and Positive Valence Systems Scale (PVSS), which provides overall, anticipatory, and consummatory anhedonia scores. PC1 scores—whose loadings indicated broad inflammatory activity, including cytokines, chemokines, and tissue remodeling/angiogenic markers—moderated the relationship between basal ganglia dopamine availability and anhedonia: At higher PC1 levels, lower dopamine availability was linked to greater overall anhedonia (as measured by SHAPS) and anticipatory anhedonia, but not to consummatory anhedonia. Exploratory analyses showed comparable effect patterns for PC4, reflecting T cell-related effector/apoptotic signaling. Analyses of individual basal ganglia regions suggest potential differences in dopaminergic-immune interactions across these regions, with effects most consistently observed in the putamen. Findings highlight nuanced neuroimmune pathways underlying anhedonia components in youth and support inflammation-based models of depression emphasizing dopaminergic-immune interplay.

## Introduction

Depression is a major public health concern given its high prevalence, recurrence, and status as a leading cause of disability worldwide (1,2). Despite this burden, approximately 30–50% of individuals respond poorly to available treatments, including antidepressant medications (3,4). Youth, in particular, face an increased risk for first-onset depression and are in a vulnerable period of neural and immune system development (5,6). Depression beginning in youth often persists into adulthood, conferring greater risk for recurrence and severe functional impairment (7,8). A symptom that often first emerges during this period—and that predicts a worse clinical course, suicidality, treatment resistance, and greater severity of depression—is anhedonia (9,10), defined as a lack of pleasure or reactivity to pleasurable stimuli (11). As a key factor in the nature and course of depression and related psychopathology, anhedonia has remained a substantial clinical challenge despite growing attention and research effort over the last five decades (12). As such, it is essential to deepen our understanding of the underlying mechanisms that remain inadequately considered in youth, a vulnerable population.

One valuable approach toward understanding the development and mechanisms of anhedonia is to adopt a more granular conceptualization that distinguishes between components potentially driven by distinct mechanisms. Notably, anticipatory and consummatory anhedonia represent different temporal phases of reactivity to pleasurable stimuli: the motivational drive or ‘wanting’ of rewards, and the experience of pleasure or ‘liking’ during reward receipt (13,14). These components involve dissociable neural pathways. Dopaminergic signaling in mesolimbic pathways is more closely linked to anticipatory as opposed to consummatory pleasure (15,16), whereas opioidergic activity has been more strongly associated with consummatory pleasure, with evidence showing that pleasurable experiences are more sensitive to mu-opioid receptor expression and function than to the dopaminergic pathway (17–19). The Research Domain Criteria (RDoC) framework situates anticipatory and consummatory components of anhedonia within the Positive Valence Systems (PVS) domain. PVS processes encompass approach motivation and responsiveness to reward cues and to reward outcomes, and anticipatory and consummatory anhedonia largely map onto these cue- and outcome-related processes. In addition, the PVS domain also includes broader reward-related constructs such as reward valuation and reward learning (11). When conceptualizing overall anhedonia within the PVS framework, it can be understood as reflecting general functioning across the system as a whole.

Beyond parsing anhedonia into components, psychoneuroimmunology models highlight the importance of conceptualizing immune activation as a biological context that shapes the impact of dopamine-mediated reward processes on anhedonia (20,21). These models are often framed in mediational terms, whereby immune activation alters dopaminergic signaling—such as synthesis, receptor binding, or postsynaptic signaling— which in turn contribute to anticipatory anhedonia (21–24). However, when considering dopamine availability rather than dopamine signaling per se, immune activation may instead be conceptualized as a moderator that amplifies how lower dopamine levels translate into motivational deficits. Mechanistically, lower dopamine availability in dopaminergic receptor–rich regions such as the basal ganglia may reduce phasic dopaminergic bursts necessary for reinforcing motivated behavior and coding reward salience. Peripheral immune activation may further reduce efficiency of central dopaminergic signaling, for example, via diminished postsynaptic sensitivity, receptor availability, and receptor binding (see Treadway et al. (24) for a comprehensive review), such that, even at comparable levels of available dopamine, downstream reward-related signaling is less effective. Under these conditions, lower dopamine availability may have a stronger impact on motivational deficits. In addition, dopaminergic reward circuits can modulate peripheral immune processes (23,25,26), suggesting a bidirectional connection that may contribute to sustained alterations across systems over time. Together, when lower dopamine availability co-occurs with immune activation, greater anticipatory anhedonia would be expected, consistent with a moderation framework.

In addition to the biological plausibility, the proposed effects are consistent with several functional magnetic resonance imaging studies examining the links between the interaction of inflammation and dopamine-mediated reward-related brain activity and anhedonia. An observational study found that higher inflammatory composite scores amplified the association between corticostriatal functional connectivity and anhedonia severity (27). Several double-blind, randomized, clinical trials on anhedonia (28,28,29) considered both patients’ levels of inflammation and reward-related neural function. Together, these studies demonstrated improvement in motivation-related and anhedonia symptoms in patients with depression and inflammation following interventions targeting dopaminergic function or inflammation (i.e., levodopa, infliximab; (28,28,29)). Although these intervention studies did not measure dopamine availability, the relevant mechanism of change requires an increase in central dopamine availability, thereby supporting the proposed interactive role of dopamine availability and inflammation.

Collectively, the mechanistic rationale and empirical research converge to support immune-dopamine interactions in anhedonia. However, most existing research on these neuroimmune processes have been conducted in adults. This leaves our understanding unclear on whether the interaction effects would similarly be observed in youth, an under-examined population yet characterized by high risk for anhedonia and developing reward-related brain and immune systems that are susceptible to changes in neuroimmune processes. A methodological challenge to execute this research is that, historically, positron emission tomography (PET) was the primary method for assessing dopamine availability. The high cost and health risks (due to radiation exposure) of PET constrain its use in youth populations.

More recently, MRI-based measures of tissue iron have emerged as validated, non-invasive proxies of dopamine levels during development, based on iron’s role in dopamine synthesis (30). Iron serves as a co-factor for tyrosine hydroxylase, the rate-limiting enzyme in dopamine synthesis, and is enriched in dopamine-rich regions such as the basal ganglia. Because iron is paramagnetic, it can be quantified using MRI-based relaxation measures, including R2’, the reversible transverse relaxation rate (reflecting the difference between R2* and R2). R2’ has been shown to quantify striatal tissue iron concentration (31,32). Importantly, tissue iron increases across adolescence (33,34), paralleling maturation of dopaminergic systems and reward-related behavior, with prior work demonstrating sufficient accumulation during youth to reliably detect variability in tissue iron (30), making this approach well-suited for developmental samples. Given developmental increases in tissue iron, age is commonly considered in analyses using R2’ measures. Tissue iron can reflect regional differences in dopamine synthesis capacity, and not for other aspects of dopaminergic function including phasic dopamine release at the synapse and receptor binding potential (30). Leveraging this non-invasive measure can help determine whether the putative reward–related neuroimmune mechanisms apply to youth. Some evidence suggests sex differences in dopaminergic processes during adolescence (35–37), although findings are mixed (30).

This study examined the moderating role of peripheral immune function in the link between dopamine availability within the basal ganglia and components of anhedonia among a single cohort of youth with current depression. Drawing on existing conceptual and empirical research, both immune and dopaminergic processes are putatively relevant to anticipatory anhedonia. Furthermore, because immune activation can impair dopaminergic efficiency, we expected that greater levels of immune activation would strengthen the association between lower dopamine availability and greater anticipatory, but not consummatory, anhedonia. Effects on overall anhedonia were additionally examined using two measures, one reflecting the classical conceptualization of anhedonia as diminished hedonic capacity (Snaith–Hamilton Pleasure Scale; (38)), and the other operationalizing anhedonia in terms of functioning within the Positive Valence Systems domain—specifically responsiveness and valuation of rewards (Positive Valence Systems Scale (11,39–41)). Overall anhedonia was indexed using total scores from both measures, whereas anticipatory and consummatory anhedonia were assessed specifically using the two PVSS subscales, which corresponds to anticipation of potential rewards and responsiveness to reward receipt, respectively (41). Findings from this study can advance the understanding of the interaction effects of immune and reward-related neural function, informing shared and distinct pathways underlying anhedonia component, and ultimately novel, integrative, and precise intervention targets among youth.

## Methods

### Participants

Fifty-five youth with depression (15–25 years; *M_age_*=21.3 years, *SD*=2.18) from the Pittsburgh area completed this study as part of the larger project, Anhedonia, Development, and Emotions: Phenotyping and Therapeutics (ADEPT), which investigated phenotyping and treatment of anhedonic depression in youth. Sample size was determined by the number of participants with relevant data from the parent study.

All participants had current depressive disorder (i.e., meeting diagnostic criteria through the Structured Clinical Interview for DSM-5 (SCID-5 (42)) and scored 12 or above on the Montgomery-Åsberg Depression Rating Scale (MADRS). Anhedonia severity (measured by Snaith–Hamilton Pleasure Scale) varied, with moderate-severe anhedonia oversampled (*M*=35.11, *SD*=6.19, *Range*=25–50). Participants were free of lifetime psychosis, bipolar disorder, moderate to severe substance use disorder over the past six months; serious, unstable cardiovascular or neurological disorders; inflammatory disorders; history of brain injury with loss of consciousness; current acute infection; past-month illicit stimulant use; pregnancy; and MRI contraindications (e.g., metal implants). Body mass index (BMI) was not an exclusionary criterion; however, physical limitations of the MRI scanner used in the larger study imposed practical constraints on body size/shape. Pharmacotherapy and psychotherapy were allowed.

All procedures were approved a priori by the University of Pittsburgh Pitt Protocol Review Online (PittPRO), and written informed consent was obtained from all participants (with assent from those under 18 years old and consent from a parent or guardian).

### Measures

#### Snaith-Hamilton Pleasure Scale (SHAPS)

The SHAPS is a self-report measure that assesses anhedonia severity (as defined by hedonic capacity (38)). This measure follows a 4-point Likert scale, with higher score reflecting more severe overall anhedonia.

#### Positive Valence System Scale

The Positive Valence System Scale (PVSS) is a self-report measure designed to assess functioning of the RDoC Positive Valence System (PVS) constructs, that is, Reward Responsiveness and Reward Valuation (41), contrasting with the SHAPS, which primarily is intended to capture the consummatory aspect of Reward Responsiveness (38). Anticipatory and consummatory anhedonia were measured using the Reward Anticipation and Initial Responsiveness subscales, respectively. The PVSS scales use a 9-point Likert scale, with lower scores reflecting more severe anhedonia. In this sample, the PVSS total, anticipatory anhedonia, and consummatory anhedonia scores were moderately associated with the SHAPS score (i.e., *r_total_*=-.63, *r_anticipatory_*=-.62, and *r_consummatory_*=-.65, respectively; Table 1).

**Table 1.**
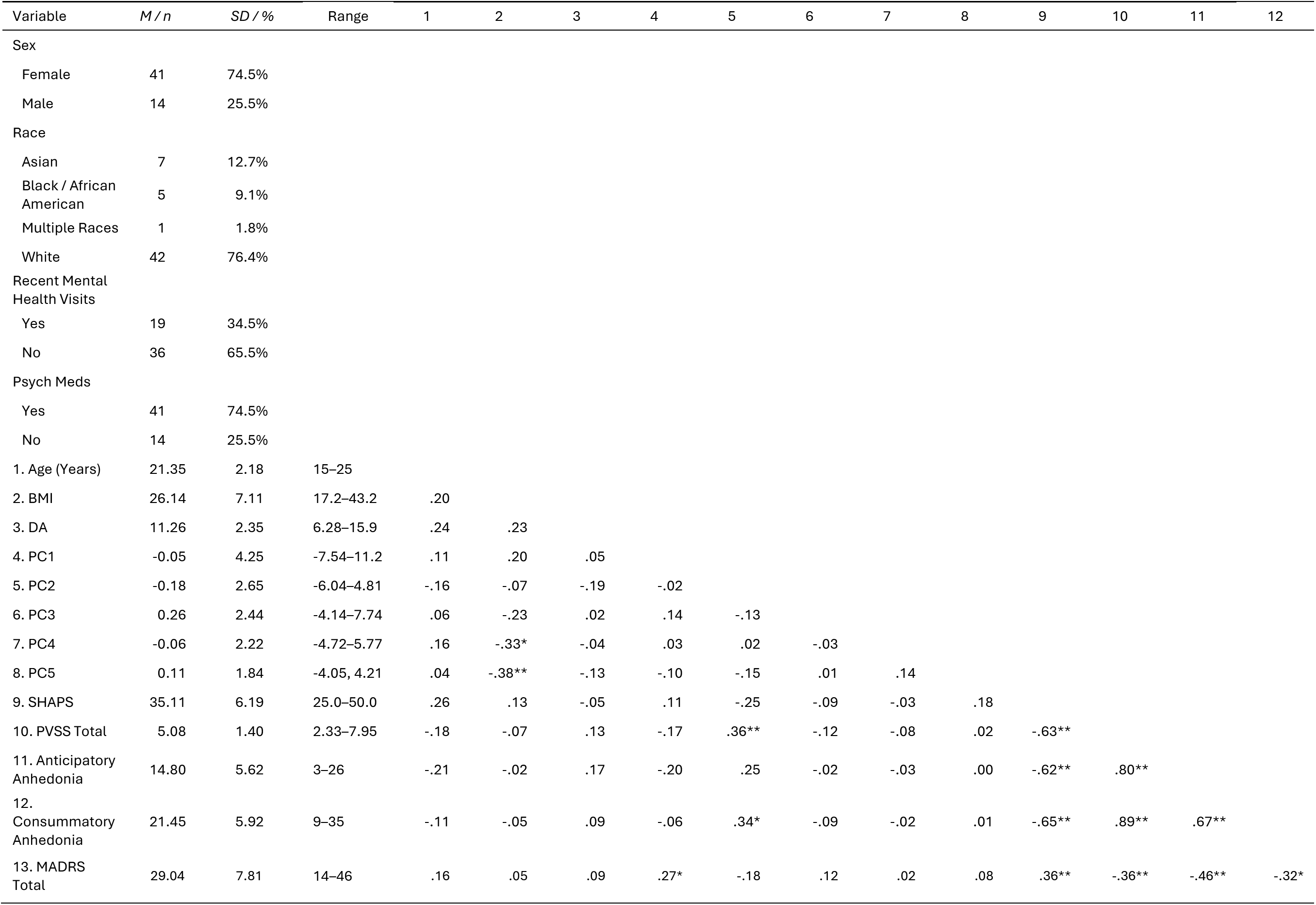
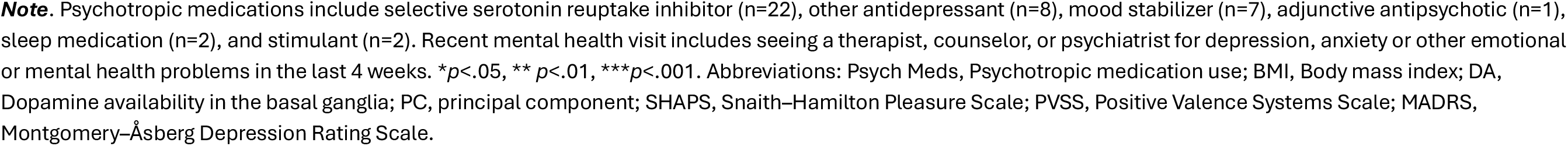
Sample characteristics and bivariate correlations of study variables (N = 55).

| Variable | M / n | SD / % | Range | 1 | 2 | 3 | 4 | 5 | 6 | 7 | 8 | 9 | 10 | 11 | 12 |
| --- | --- | --- | --- | --- | --- | --- | --- | --- | --- | --- | --- | --- | --- | --- | --- |
| Sex |  |  |  |  |  |  |  |  |  |  |  |  |  |  |  |
| Female | 41 | 74.5% |  |  |  |  |  |  |  |  |  |  |  |  |  |
| Male | 14 | 25.5% |  |  |  |  |  |  |  |  |  |  |  |  |  |
| Race |  |  |  |  |  |  |  |  |  |  |  |  |  |  |  |
| Asian | 7 | 12.7% |  |  |  |  |  |  |  |  |  |  |  |  |  |
| Black / African American | 5 | 9.1% |  |  |  |  |  |  |  |  |  |  |  |  |  |
| Multiple Races | 1 | 1.8% |  |  |  |  |  |  |  |  |  |  |  |  |  |
| White | 42 | 76.4% |  |  |  |  |  |  |  |  |  |  |  |  |  |
| Recent Mental Health Visits |  |  |  |  |  |  |  |  |  |  |  |  |  |  |  |
| Yes | 19 | 34.5% |  |  |  |  |  |  |  |  |  |  |  |  |  |
| No | 36 | 65.5% |  |  |  |  |  |  |  |  |  |  |  |  |  |
| Psych Meds |  |  |  |  |  |  |  |  |  |  |  |  |  |  |  |
| Yes | 41 | 74.5% |  |  |  |  |  |  |  |  |  |  |  |  |  |
| No | 14 | 25.5% |  |  |  |  |  |  |  |  |  |  |  |  |  |
| 1. Age (Years) | 21.35 | 2.18 | 15–25 |  |  |  |  |  |  |  |  |  |  |  |  |
| 2. BMI | 26.14 | 7.11 | 17.2–43.2 | .20 |  |  |  |  |  |  |  |  |  |  |  |
| 3. DA | 11.26 | 2.35 | 6.28–15.9 | .24 | .23 |  |  |  |  |  |  |  |  |  |  |
| 4. PC1 | -0.05 | 4.25 | -7.54–11.2 | .11 | .20 | .05 |  |  |  |  |  |  |  |  |  |
| 5. PC2 | -0.18 | 2.65 | -6.04–4.81 | -.16 | -.07 | -.19 | -.02 |  |  |  |  |  |  |  |  |
| 6. PC3 | 0.26 | 2.44 | -4.14–7.74 | .06 | -.23 | .02 | .14 | -.13 |  |  |  |  |  |  |  |
| 7. PC4 | -0.06 | 2.22 | -4.72–5.77 | .16 | -.33* | -.04 | .03 | .02 | -.03 |  |  |  |  |  |  |
| 8. PC5 | 0.11 | 1.84 | -4.05, 4.21 | .04 | -.38** | -.13 | -.10 | -.15 | .01 | .14 |  |  |  |  |  |
| 9. SHAPS | 35.11 | 6.19 | 25.0–50.0 | .26 | .13 | -.05 | .11 | -.25 | -.09 | -.03 | .18 |  |  |  |  |
| 10. PVSS Total | 5.08 | 1.40 | 2.33–7.95 | -.18 | -.07 | .13 | -.17 | .36** | -.12 | -.08 | .02 | -.63** |  |  |  |
| 11. Anticipatory Anhedonia | 14.80 | 5.62 | 3–26 | -.21 | -.02 | .17 | -.20 | .25 | -.02 | -.03 | .00 | -.62** | .80** |  |  |
| 12. Consummatory Anhedonia | 21.45 | 5.92 | 9–35 | -.11 | -.05 | .09 | -.06 | .34* | -.09 | -.02 | .01 | -.65** | .89** | .67** |  |
| 13. MADRS Total | 29.04 | 7.81 | 14–46 | .16 | .05 | .09 | .27* | -.18 | .12 | .02 | .08 | .36** | -.36** | -.46** | -.32* |

**Note.** Psychotropic medications include selective serotonin reuptake inhibitor (n=22), other antidepressant (n=8), mood stabilizer (n=7), adjunctive antipsychotic (n=1), sleep medication (n=2), and stimulant (n=2). Recent mental health visit includes seeing a therapist, counselor, or psychiatrist for depression, anxiety or other emotional or mental health problems in the last 4 weeks. \* $p < .05$ , \*\* $p < .01$ , \*\*\* $p < .001$ . Abbreviations: Psych Meds, Psychotropic medication use; BMI, Body mass index; DA, Dopamine availability in the basal ganglia; PC, principal component; SHAPS, Snaith–Hamilton Pleasure Scale; PVSS, Positive Valence Systems Scale; MADRS, Montgomery–Åsberg Depression Rating Scale.

#### The Structured Clinical Interview for DSM-5 (SCID-5)

The SCID-5 (42) is a semi-structured clinical interview designed to assess current and past diagnostic history of psychopathology. This interview was conducted by trained ADEPT research assistants and postdoctoral fellows.

#### Montgomery-Åsberg Depression Rating Scale (MADRS)

The MADRS (43) is a 10-item interviewer-rated questionnaire assessing depressive symptoms. This measure used a 7-point Likert scale, with higher scores indicating more severe depression.

#### Dopamine Availability in the Basal Ganglia

R2’, a quantitative MR-based measure (44), was used to measure basal ganglia tissue iron as a proxy of dopamine availability (30). Prior work integrating PET and quantitative MRI has demonstrated that R2’ is moderately and consistently associated with presynaptic dopamine levels, particularly in the ventral striatum, while the strength may vary across basal ganglia subregions (30). R2’ reflects the reversible transverse relaxation rate (1/T2’), calculated across the basal ganglia and within each of the subregions (i.e., caudate, putamen, pallidum, and nucleus accumbens [NAc]; see Figure 1 for masks using Harvard-Oxford Atlas) as the difference between the effective R2* (1/T2*) and irreversible R2 (1/T2) relaxation rates (44).

**Figure 1.**
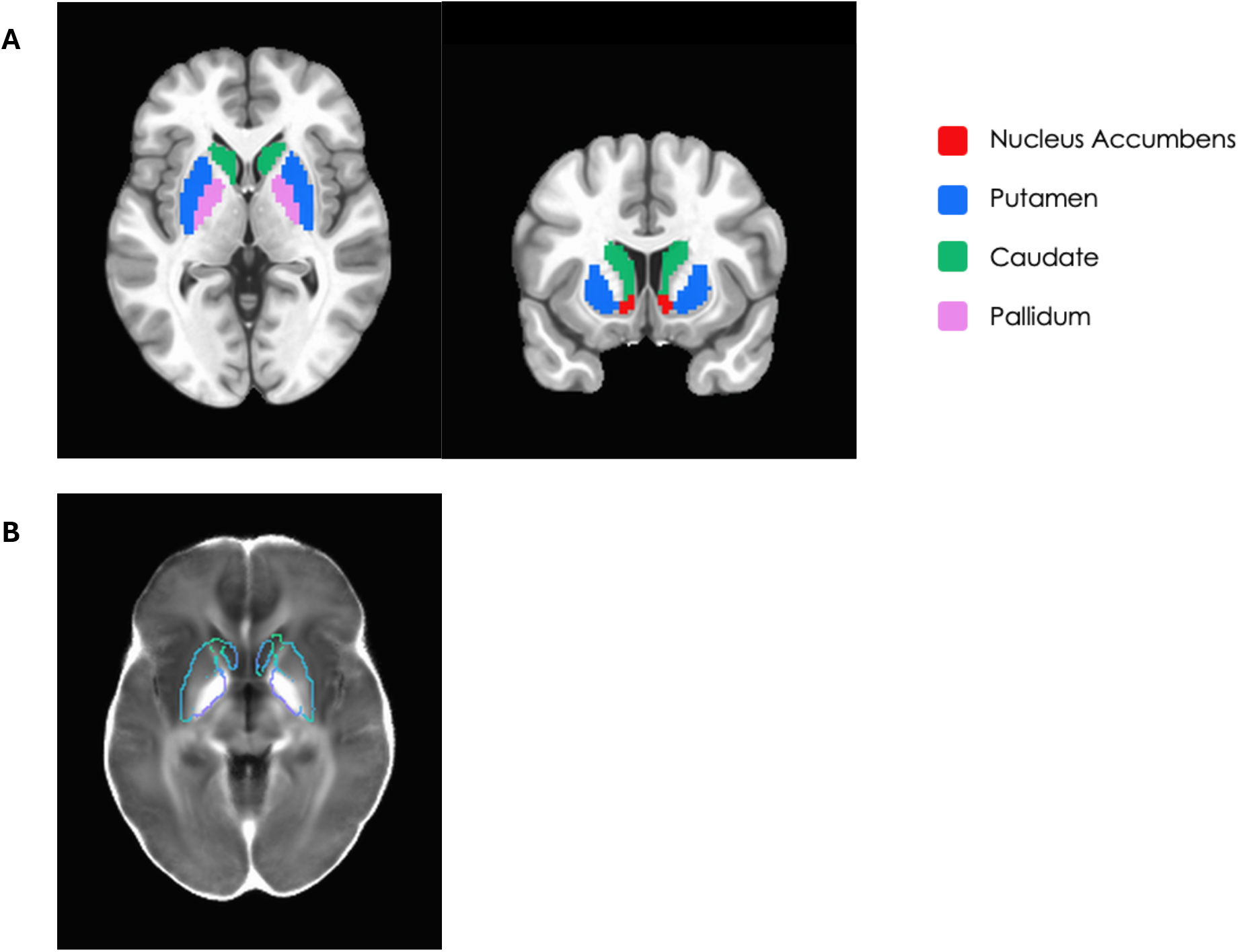
Masks and mean R2’ signals within basal ganglia regions. The figures illustrate **(A)** masks of basal ganglia in axial (left) and coronal (right) views, including nucleus accumbens (red), putamen (blue), caudate (green), pallidum (pink), with basal ganglia defined as the union of these four subregions; and **(B)** mean R2’ signal within basal ganglia, averaged across participants passing quality control. Brighter intensity reflects higher R2’ values, indicative of greater tissue iron. Atlas-based masks of basal ganglia subregions are overlaid for anatomical reference.

Following previous work by our group (30), R2* data were acquired using a multi-echo gradient echo (mGRE) sequence (TE=3.8, 8, 18, 23 ms; TR=1110 ms, flip angle=25°, FoV=240×240 mm, 40 slices with 0.938×0.938 mm in place resolution and 3 mm spacing). R2 data were acquired using a multi-echo turbo spin echo (mTSE) sequence (TE=12, 99, 186 ms, TR=10800 ms, FoV=240×240 mm, 40 slices with 0.938×0.938 mm in place resolution and 3 mm spacing). R2 and R2* were estimated using a quadratic penalized least squares (QPLS) method (following Funai et al., IEEE Trans. Med. Imaging, 2008) to fit multi-echo data to a mono-exponential decay model (see Larsen et al. (30) for full description). R2* and R2 images were registered to MNI space using AFNI (Cox, 1996), using an affine registration to the anatomical, and non-linear warp to MNI space based on the registration of the anatomical to the MNI template. Estimates of R2’ were derived by subtracting R2 from R2* estimates. All R2’ images were visually assessed by trained research personnel for motion, shimming, susceptibility, or registration artifacts. Scans with visually identifiable artifacts were excluded from all iron analyses (*n*=15).

Given the putative functional distinctions among basal ganglia subregions (i.e., caudate, putamen, pallidum, and NAc), secondary analyses were conducted in which each subregion replaced the combined basal ganglia signal for any effects observed at the whole–basal ganglia level.

#### Assay for Immune-Related Proteins

Blood was collected into a BD vacutainer tube via venipuncture on the same day as the R2’ scan (9 a.m.– 5 p.m.). To avoid confounding effects, participants were instructed to avoid alcoholic beverage overnight prior to the blood draw. Thirty minutes later, the collected samples were centrifuged (15 minutes at 3000 RPM) to separate plasma, which were aliquoted and then stored in a −80°C freezer until the time of analysis. Plasma samples were assayed and analyzed for levels of immune-related proteins, using Olink Target 96 Inflammation panel (Olink Bioscience, Uppsala, Sweden), according to the manufacturer’s instructions, at the Human Immune Monitoring Center of the Icahn School of Medicine at Mount Sinai. This panel measures 92 target proteins (principally chemokines, interleukins, growth factors, TNF-related molecules, other inflammation-related proteins). The assay provides relative quantification of these targets, expressed as normalized protein eXpression (NPX) unit on a log2 scale. Higher NPX value indicates higher protein concentration. Following established procedures, if a specific target in the panel yielded values for which ≥25% of the samples were lower than the limit of detection or did not pass quality control (based on internal controls within samples [within ±0.3 NPX of the median] and variability of controls across plates [SD<0.2 NPX across controls]), the target overall was excluded from the analyses. This resulted in 70 immune proteins included for further analyses (see Supplementary Table 1 for descriptive statistics of included proteins and Supplementary List of Abbreviations for full protein names).

### Principal Component Analysis (PCA) for Immune Proteins

To reduce dimensionality of the immune protein data, PCA was conducted following prior work (45). Individual outliers (i.e., SD >3 from the group mean; prevalence listed in Supplementary Table 2) were treated as missing and imputed using a principal component-based method (imputePCA from missMDA R package), which estimates values based on the underlying low-dimensional structure of the data. The optimal number of principal components to retain was determined using a combination of criteria, including inspection of variance explained and Horn’s parallel analysis (46), where components are retained when their eigenvalues exceed those derived from randomly simulated data. Both approaches indicate inflection after the fifth component (details in Results). Accordingly, five principal components were retained, collectively explaining approximately 55% of the total variance. To optimize interpretability, an oblimin rotation was applied. We were primarily interested in the component reflecting general immune activation, and its corresponding component score (i.e., each participant’s value on the component), computed as weighted linear combination of the original standardized markers, was used for primary analyses. Exploratory analyses were conducted using the remaining components. Unrotated PCA solution was also applied to assess robustness. In sensitivity analyses, outlier values were excluded rather than imputed (see Supplementary Materials for details).

### Statistical Analyses

Multiple linear regressions examined interaction effects between basal ganglia dopamine availability (independent variable) and immune PC scores (moderators) on overall, anticipatory, and consummatory anhedonia severity (dependent variables). Significant interactions were probed using the Johnson-Neyman procedure to identify ranges of immune PC scores at which dopamine availability was associated with anhedonia. Secondary analyses probed effects within basal ganglia subregions (i.e., caudate, NAc, pallidum, putamen). Exploratory analyses repeated the above models using the remaining retained immune PCs, and sensitivity analyses excluding rather than imputing outlier values. Additional analyses assessed robustness using the unrotated solution. Covariates were sex assigned at birth, BMI, age, and current psychotropic medication use (yes/no; e.g., antidepressants, mood stabilizers, antipsychotics, sleep medications, benzodiazepines), selected based on known associations with immune and/or brain functions (47–49). Models additionally adjusted for depressive symptom severity (excluding anhedonia) to isolate effects specific to anhedonia. Attention-deficit/hyperactivity disorder (ADHD) status was included in separate analyses to evaluate robustness of findings. Sample characteristics and correlations among study variables are displayed in Table 1.

### Code Availability

R code used for analyses is available upon reasonable request.

## Results

### Preliminary Analyses

#### PCA Results: Immune Activation

Figure 2 illustrates variance explained by principal components the 70 immune-related proteins. Consistent with parallel analysis, five principal components were retained, accounting for 55% of total variance, with individual components explaining 24%, 10%, 8%, 8%, and 5% respectively. Rotated and unrotated loadings for the 70 immune proteins are provided in Supplementary Tables 3 and 4. Scores from these retained principal components (i.e., weighted linear combinations of the immune markers) were extracted to represent each participant’s expression of the corresponding components.

**Fig. 2:**
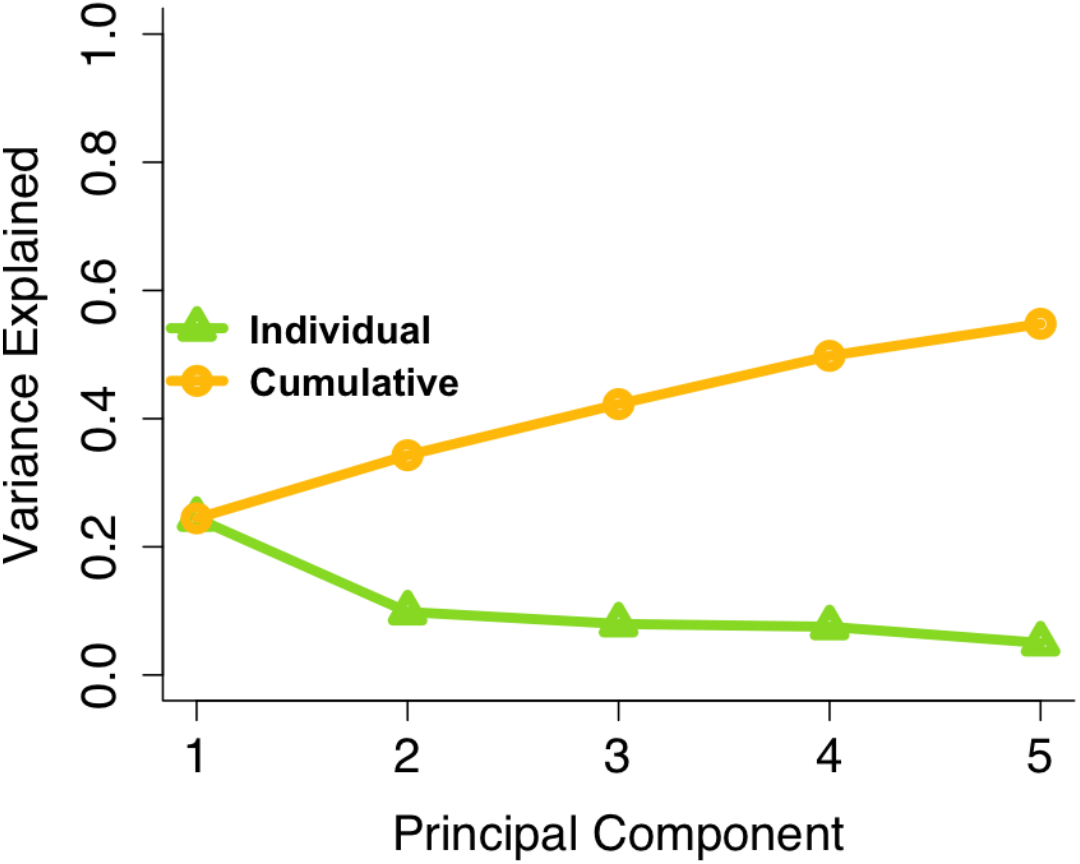
Principal Component Analysis Results: Immune Activation. The plot illustrates the variance explained by principal component derived from 70 immune-related proteins. Based on parallel analysis, five principal components were retained, accounting for 55% of the total variance, with individual components explaining 24%, 10%, 8%, 8% and 5% respectively.

Following oblimin rotation, the first principal component (PC1) was characterized by high positive loadings on markers including pro-inflammatory cytokines (e.g., IL-6, IL-8), chemokines involved in immune cell recruitment (e.g., MCP-1, MCP-2, MCP-4), and markers of tissue remodeling and angiogenesis (e.g., VEGFA, HGF), consistent with coordinated inflammatory profile—the construct of primary interest in this study. Markers with the highest loadings on PC1 were predominantly associated with the myeloid lineage (e.g., monocytes/macrophage-derived proteins), suggesting that this component reflects variation in innate immune activity. Thus, primary analyses focused on PC1, which showed a comparable pattern across rotated and unrotated solutions. Exploratory analyses examined the remaining components (PC2-PC5).

### Primary Analyses

#### General Immune Activation (PC1) and Anhedonia

Main effects were found for higher immune activation (PC1 score) on more severe SHAPS overall anhedonia (*B*=19.561, *SE*=6.122, *p*=.003, *ΔR^2^*=.142) and PVSS anticipatory anhedonia (*B*=-13.344, *SE*=5.348, *p*=.016, *ΔR^2^*=.080). Immune activation was unrelated to PVSS overall anhedonia (*B*=-.115, *SE*=.181, *p*=.527, *ΔR^2^*=.007) and PVSS consummatory anhedonia (*B*=-.292, *SE*=.812, *p*=.721, *ΔR^2^*=.002).

#### Basal Ganglia Dopamine Availability and Anhedonia

We did not detect main effects for dopamine availability within the basal ganglia on SHAPS overall anhedonia (*B*=-.584, SE=.339, *p*=.092, *ΔR^2^*=.041), PVSS overall anhedonia (*B*=.115, *SE*=.081, *p*=.165, *ΔR^2^*=.033) or PVSS consummatory anhedonia (*B*=.391, *SE*=.365, *p*=.290, *ΔR^2^*=.021), except for PVSS anticipatory anhedonia (*B*=.761, *SE*=.296, *p*=.013, *ΔR^2^*=.085).

#### Interaction of Basal Ganglia Dopamine Availability and Immune Activation in Relation to Anhedonia

There was an interaction effect between dopamine availability in the basal ganglia and immune activation on SHAPS overall anhedonia (*B*=-1.715, *SE*=.522, *p*=.002, *ΔR^2^=*.150), such that for participants in the top 63.79% of immune activation, lower dopamine availability was associated with more severe SHAPS overall anhedonia (Figure 3A). The significant region at the lower end of PC1 values (Figure 3A) represented only a small portion of observed sample distribution (n=8) and should therefore be interpreted with caution. No interaction effect was detectd for PVSS overall anhedonia (*B*=.208, *SE*=.126, *p*=.107, *ΔR^2^*=.043; Figure 3B).

**Fig. 3.**
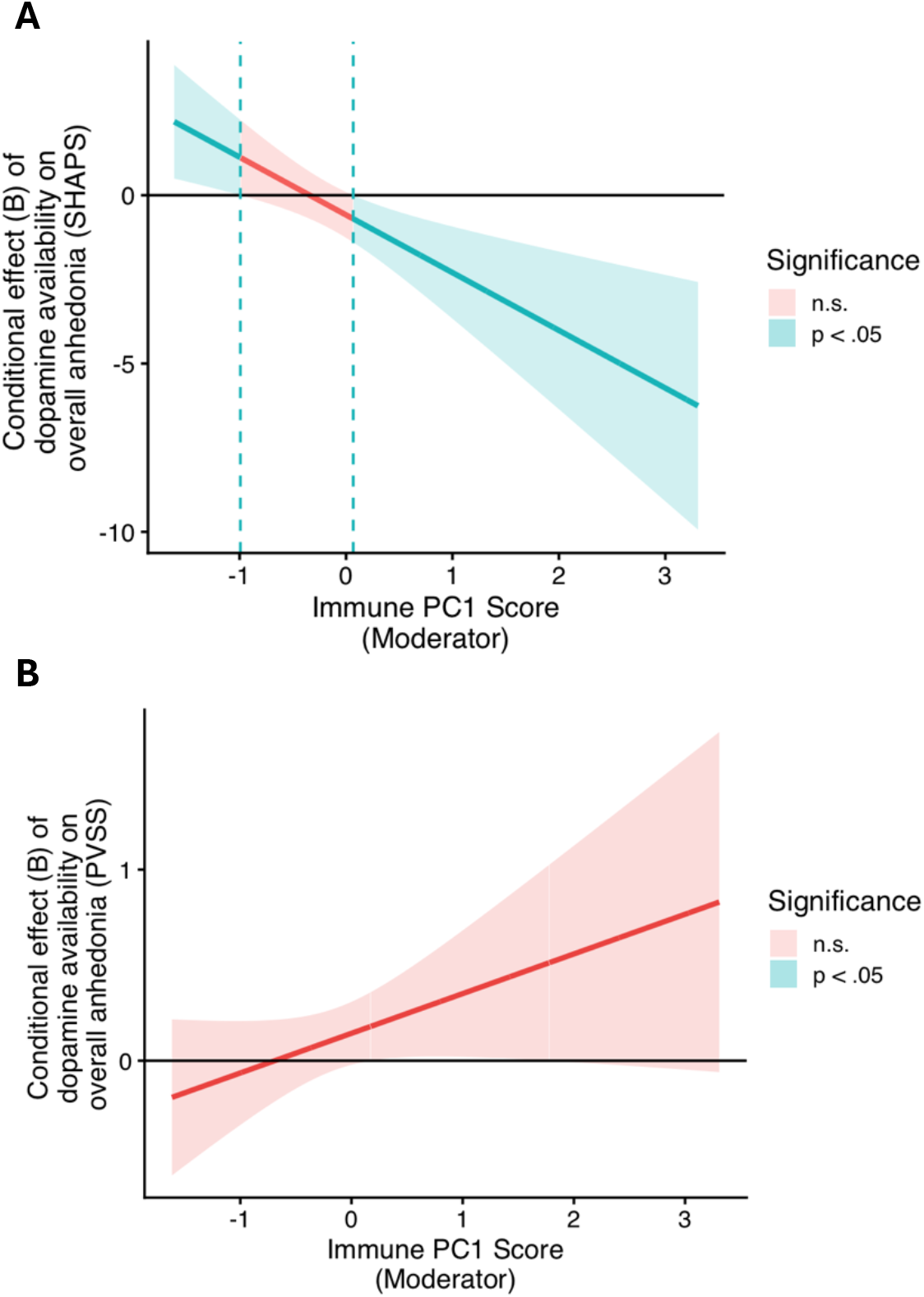
Conditional effects of dopamine availability in the basal ganglia on overall anhedonia by levels of general immune activation. Johnson-Neyman plots showing the magnitude of association between dopamine availability in the basal ganglia and overall anhedonia (y-axis), as a function of general immune activation (Principal Component 1; x-axis). Overall anhedonia was assessed using the Snaith–Hamilton Pleasure Scale (SHAPS; A) and the Positive Valence Systems Scale (PVSS; B). Shaded regions indicate values of the moderator (Principal Component 1) at which the conditional effect of dopamine availability on overall anhedonia differs significantly from zero (*p* <.05; blue) versus not significant (n.s.; *p* ≥ .05; red). The solid line represents the estimated conditional effect, and shaded bands denotes 95% confidence intervals. Estimates are shown across the observed range of immune Principal Component 1 values in the sample. **A.** At higher levels of general immune activation, lower dopamine availability in the basal ganglia was associated with greater overall anhedonia (as measured by the SHAPS). The significant region at the lower end of PC1 values (i.e., PC1 values <.99; blue) reflects a relatively small portion of the observed distribution (approximately the bottom 13.79% of values; n=8) and should be interpreted with caution. **B.** The moderation effect was not observed when overall anhedonia was measured by the PVSS.

There was also an interaction effect for PVSS anticipatory anhedonia (B=1.123, *SE*=.456, *p*=.018, *ΔR^2^*=.078), such that for participants in the top 53.45% of immune activation, lower dopamine availability was associated with more severe PVSS anticipatory anhedonia (Figure 4A). No interaction effect was detected for PVSS consummatory anhedonia (*B*=.767, *SE*=.571, *p*=.186, *ΔR^2^*=.033; Figure 4B).

**Fig. 4.**
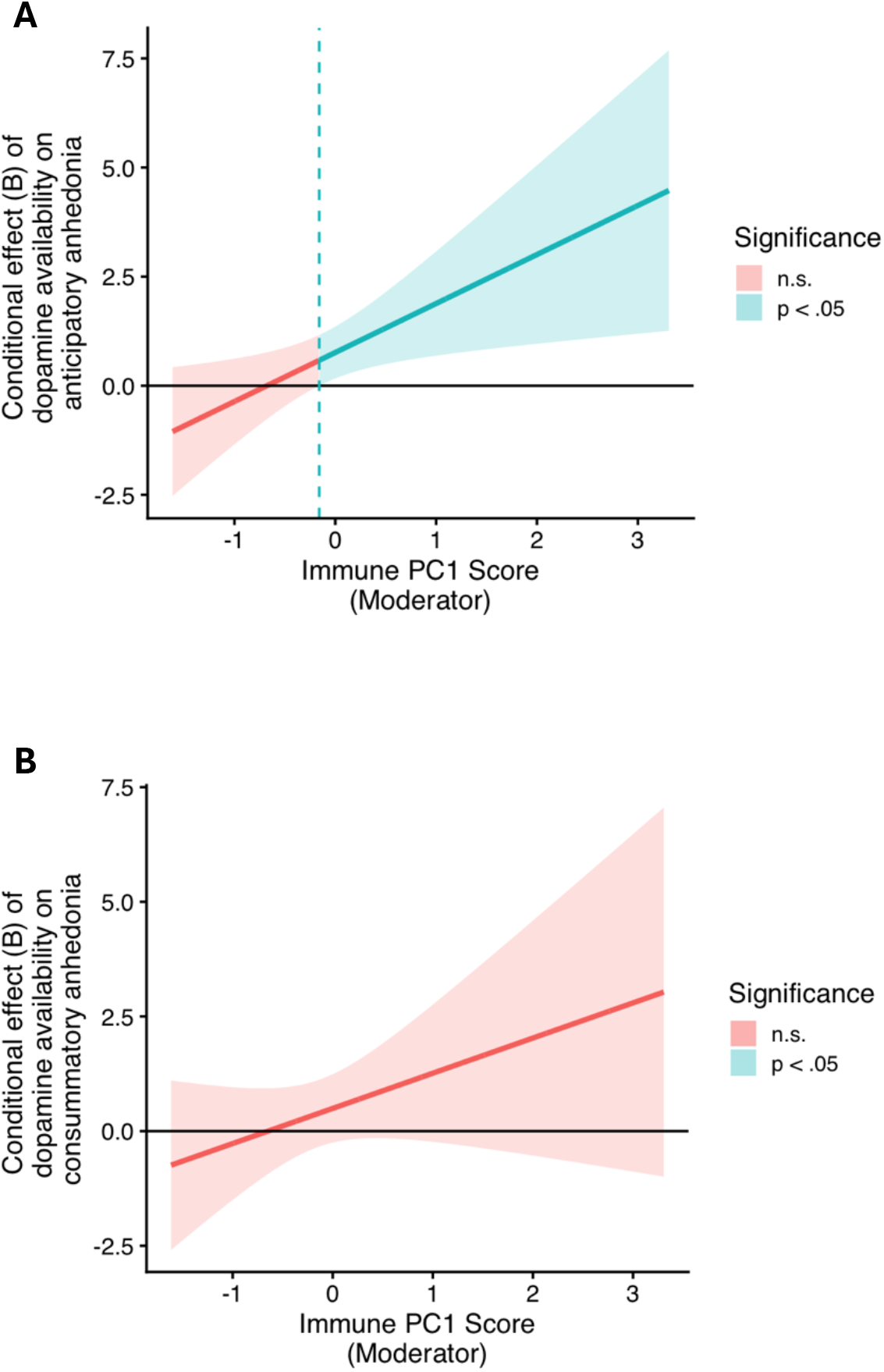
Conditional effects of dopamine availability in the basal ganglia on anhedonia components by levels of general immune activation. Johnson-Neyman plots showing the magnitude of association between dopamine availability in the basal ganglia and components of anhedonia (i.e., anticipatory anhedonia [A] and consummatory anhedonia [B] (y-axis); as measured by the subscales of Positive Valence Systems Scale), as a function of general immune activation (Principal Component 1; x-axis). Shaded regions indicate values of the moderator (Princiapl Component 1) at which the conditional effect of dopamine availability on anhedonia component differs significantly from zero (*p* <.05; blue) versus not significant (n.s.; *p* ≥ .05; red). The solid line represents the estimated conditional effect, and shaded bands denotes 95% confidence intervals. Estimates are shown across the observed range of immune Principal Component 1 values in the sample. **A.** At higher levels of general immune activation, lower dopamine availability in the basal ganglia was associated with greater anticipatory anhedonia. **B.** The moderation effect was not observed for consummatory anhedonia.

### Secondary and Exploratory Analyses

Secondary analyses probed effects within basal ganglia subregions for significant findings identified in the primary analyses, and exploratory analyses examined associations with the four other immune PC scores. Dopamine Availability X Immune PC1 interaction effects were present for several basal ganglia subregions for SHAPS overall anhedonia (caudate, pallidum, putamen) and anticipatory anhedonia (putamen). Full results are reported in Supplementary Material.

There was a Dopamine Availability X Immune PC4 interaction effect on SHAPS overall anhedonia and PVSS anticipatory anhedonia. For participants in the top 60.34% and 37.93% of PC4 scores, respectively, lower dopamine availability in the basal ganglia was associated with greater SHAPS overall anhedonia and PVSS anticipatory anhedonia. Follow-up analyses indicated effects within the putamen and pallidum. There were no effects with the remaining PC scores.

### Sensitivity and Robustness

Full sensitivity and robustness results are reported in the Supplementary Material. Sensitivity analyses for the primary analyses (interaction effects between basal ganglia dopamine availability and Immune PC1 on anhedonia outcomes) using a complete-case approach (i.e., excluding rather than imputing outlier values) showed effect in the same direction as the primary analyses, although statistical significance was attenuated, likely reflecting reduced sample size (N=35) and corresponding power.

Analyses were repeated using the unrotated PC solution, and with ADHD status as an additional covariate. Findings were directionally consistent with the primary analyses, indicating that results were not driven by the rotation method or ADHD status.

## Discussion

This study examined whether peripheral immune processes and naturally occurring dopamine availability in the basal ganglia, alone and in interaction, were associated with anhedonia among youth with current depression. An expected principal component (PC1) reflecting coordinated inflammatory activity and innate, myeloid-driven immune processes was identified. This component includes high-loading markers (e.g., IL-6, IL-8, MCPs (50–52)) that are established indicators of inflammatory activity, have been associated with depressive symptoms in prior work, and have been detectable in youth samples, including those with depression (52). As hypothesized, greater anhedonia was associated with higher immune PC1 scores. Likewise, lower dopamine availability was associated with greater anticipatory anhedonia (14). Aligned with conceptual and empirical work on immune pathways in anhedonia (20,53), general immune activation amplified the association between lower dopamine availability in the basal ganglia and greater anhedonia severity. Moreover, consistent with prior research suggesting that mechanisms of anhedonia vary by component (14,54), the associations of immune processes and dopamine availability with anhedonia differ across anhedonia components. Exploratory analyses preliminarily suggest differences among subregions in these effects, with patterns most consistently observed in putamen, although replication in larger samples is necessary to confirm these findings.

The robust amplification effects between low dopamine availiablity and general immune activation detected for greater anticipatory anhedonia as compared to consummatory anhedonia aligns with the theoretical rationale and neurobiological research suggesting that immune activation is relevant to shifting the motivational state and the function in dopaminergic pathways subserving approach motivation (21,24,55). By showing that distinct neural and immune pathways, separately and interactively, may characterize different components of anhedonia, these findings underscore the risk of aggregating anhedonia into a unitary construct without examining or conflating its components (13,14,54). Such lumping can, from an analytical standpoint, dilute true effects and obscure pathway-specific associations. From a precision medicine perspective, failing to disaggregate separate components of anhedonia when examining pathophysiology can increase the risk of mismatching mechanisms with symptom presentations and thus generating imprecise treatment targets.

Notably, interaction effects were detected for overall anhedonia when measured by the SHAPS, but not by the PVSS total, although both are expected to capture the same construct. The association between immune activation and SHAPS scores aligns with prior findings, both cross-sectionally in adolescent samples with diverse psychiatric conditions (56) and longitudinally in adolescents with depressive disorders (57). Neither study included the PVSS, however, and therefore could not determine whether the absence of effect was specific to PVSS total or reflected a broader measurement-related discrepancy. One possible explanation for this discrepancy is that the SHAPS, widely considered a gold-standard anhedonia measure, was developed based on the classical definition of anhedonia as a lack of pleasure (38) and, as a result, is often interpreted as indexing consummatory anhedonia. Yet this interpretation appears unlikely, as significant effects were observed for the SHAPS but not the PVSS consummatory subscale, which was designed to capture variability in hedonic response to recipt of reward. Moreover, SHAPS scores were moderately correlated with PVSS consummatory anhedonia (*r*=-.65) and in similar magnitude with PVSS total (*r*=-.63), suggesting that the SHAPS captures anhedonia more broadly than just consummatory anhedonia, while remaining partially dissociable from PVSS total.

A more plausible explanation for this discrepancy lies in differences in item phrasing between the measures. Although the SHAPS is often interpreted as assessing hedonic capacity over the past few days, its items are framed hypothetically (i.e., “*I would* find pleasure in…,” “*I would* enjoy…”), requiring participants to evaluate expected pleasure in imagined scenarios. This format can introduce future-oriented and conditional framing and bias responses towards anticipatory pleasure or motivational appraisal, thereby blurring anticipatory versus consummatory anhedonia assessment. In contrast, PVSS consummatory items are anchored in affective responses to recent actual experience (e.g., “I *savored* …,” “I *felt pleased* when…”), more directly capturing in-the-moment pleasure (41). Thus, SHAPS scores may reflect a broader hedonic capacity that, as an unintended consequence of its hypothetical phrasing, include anticipatory appraisals, whereas the PVSS consummatory subscale may more selectively index experienced pleasure. This distinction may help explain why SHAPS, but not the PVSS consummatory subscale, showed associations in the present analyses. Because PVSS total represents the broad construct spanning general PVS deficits beyond anticipatory and consummatory dimensions, the null result further suggests that collapsing across components may attenuate associations if some PVS components are less relevant to the processes. Such aggregation may obscure component-specific effects, underscoring the value of targeted measures that capture distinct anhedonia components in future research (11,14,58).

Our exploratory analyses identified interactive effects involving other immune components. At higher immune PC4 scores (reflecting T cell-related effector and apoptotic signaling, characterized by high positive loadings on 4E-BP1, CD8A, uPA, TRAIL, CCL11, IL-17C, TNF, and high negative loadings on CX3CL1, TNF-β), lower basal ganglia dopamine availability was associated with greater SHAPS overall anhedonia and anticipatory anhedonia. No effects were observed for immune PC2 (high positive loadings on STAMBP, SIRT2, AXIN1, ST1A1, CASP-8, CXCL1, CXCL11, CXCL5, CXCL6, CD40), PC3 (high positive loadings on CD5, CD244, CD6, CXCL10, TNFRSF9, CCL3, IFN-γ, CXCL9, CDCP1, IL10), or PC5 scores (high positive loadings on DNER, TWEAK). These findings collectively underscore that immune processes are not monolithic. Rather, distinct constellations and contrasts among immune processes (e.g., proinflammatory vs. adaptive pathways) may yield diffrent clinical implications across specific symptom dimensions. However, most prior research on immune function in relation to psychopathology has focused on elevated levels of individual markers or small cyokine panels—an approach that may overlook broader patterns of immune activity relevant to symptom heterogeneity. In contrast, our analyses leveraged a large-scale multiplex assay—an innovative approach that enabled comprehensive immune profiling across numerous markers. This approach provided substantially greater granularity than traditional single-marker methods and allowed immune pathways (e.g., inflammatory and antiviral pathways) to be examined both independently and jointly, offering a more nuanced understanding of how immune function relates to psychopathology. Given the exploratory nature of these analyses, future research deriving factors from large sets of immune markers should replicate this effort using a confirmatory analytical approach.

Given that most studies on brain-immune pathways and related interventions have been conducted in adults, a critical question has been whether such findings extend to other developmental stages, as developmental context may shape how immune, dopaminergic, and affective systems interact, and, in turn, how these interactions confer risk for anhedonia (6). For youth, this context is characterized by ongoing maturation of mesocorticolimbic dopaminergic pathways and immune signaling processes. Because these systems are highly plastic and mutually influential during this period (6), disruptions in one are likely to affect the other, thereby heightening risk for anhedonia. However, whether empirical research supports this theoretical notion remains unclear, as this area of research is generally underdeveloped in youth. The current study begins to bridge this gap, providing preliminary support for the potential generalizability of prior mechanistic and interventional findings to a younger developmental stage. Further research investigating brain-immune mechanisms underlying anhedonia within developmental contexts will be valuable for informing early, targeted, and developmentally sensitive interventions for this vulnerable age group.

This study integrated multiple levels of analysis from self-report measures to biological and neural indices, providing a comprehensive understanding of pathways underlying anhedonia. It also addressed a gap in the literature by examining interactions between two systems typically studied independently: reward-related (dopaminergic) neural function and immune activation. Most prior work has hypothesized *mediational* pathways in which immune activation impairs dopaminergic function to induce anhedonia, a question best tested in longitudinal designs, especially in youth undergoing rapid maturation. The current findings instead highlight the complementary value of an moderation model, wherein immune activation may create a biological context that renders available dopamine less effective in supporting reward-related function. Notably, this is among the first studies to apply the tissue iron technique, an advance that allows for greater insight into neurochemical mechanisms of psychopathology in youth, which is typically under-examined due to the limitations of PET and other molecular imaging approaches for younger populations. Consistent with expectations, the dopamine variable was detectable in the sample even though tissue iron is deposited during development. The cross-system focus, alongside consideration of methodological issues (e.g., lumping symptoms) may help explain inconsistencies across prior studies (59,60).

Nevertheless, several limitations should be considered when interpreting these findings. First, given the cross-sectional, observational design, results are correlational and do not establish causality. Although direct manipulation of dopaminergic function is not feasible, experimental inflammation paradigms (e.g., immune challenge) and pharmacological approaches may help strengthen causal inference. Second, while focusing on youth is a strength for understanding development and early course of anhedonia, it limits assessment of specificity and generalizability across the lifespan. Third, although tissue iron provides an indirect index of dopamine availability, other aspects of dopaminergic function (e.g., synaptic release, receptor availability) would require PET, whose elective use for research purposes is not ethically feasible in younger participants. Additionally, null effects in the NAc may reflect greater susceptibility to artifact-related signal distortion (e.g., blooming) in small subcortical regions and reduced power due to data exclusion. Future work using larger samples and complementary approaches (e.g., corticostriatal functional connectivity) may help clarify these relationships. Fourth, we could not adequately examine sex or racial differences due to sample imbalance and limited diversity, despite evidence that these factors contribute to variability in immune and affective processes (61–65). Future work in more diverse samples is needed to examine these differences, particularly in youth, whose neural, immune, and hormonal systems are undergoing rapid and dynamic maturation, alongside increasing exposure to complex environmental and social input (6). Finally, key reward processes relevant to anhedonia, particularly reward learning, were not assessed, limiting conclusions about anhedonia and reward processing alterations.

Taken together, our findings suggest that different neural and immune pathways may be uniquely associated with different components of anhedonia in youth and underscore the importance of jointly assessing reward and immune processes when studying anhedonia. This integrated approach holds promise for informing more targeted treatments for this debilitating symptom.

## Acknowledgments

The larger funded study, “Anhedonic Depression in Adolescents and Young Adults: Capturing Phenotypes and Disrupting Mechanisms at a Critical Developmental Point” was supported by Wellcome Leap as part of the Multi-Channel Psych Program. Wellcome Leap MCPsych evaluated the manuscript prior to publication for IP considerations but did not influence any of the analyses. This work was additionally funded by R01 MH127014 and R01 MH124900 (PI: EEF), T32MH015750 (IKC), the American Psychological Foundation’s Peter and Malina James & Dr. Louis P. James Legacy Scholarship (IKC), and the University of Pittsburgh Medical Center Psychiatry Research Pathway (KT). The authors would like to thank the participants and their families for participation in this study and Ashley Pogue for assistance in data collection and management.

## References

1. Murray CJL, editor. The global burden of disease: a comprehensive assessment of mortality and disability from diseases, injuries, and risk factors in 1990 and projected to 2020; summary. Cambridge: Harvard School of Public Health [u.a.]; 1996. 43 p. (Global burden of disease and injury series; no. 1).

2. Collaborators UB of D. The State of US health, 1990-2010: Burden of diseases, injuries, and risk factors. JAMA - J Am Med Assoc. 2013 Aug 14;310(6):591–608. doi:10.1001/jama.2013.13805 PubMed PMID: 23842577.

3. Rush AJ, Trivedi MH, Wisniewski SR, Nierenberg AA, Stewart JW, Warden D, et al. Acute and Longer-Term Outcomes in Depressed Outpatients Requiring One or Several Treatment Steps: A STAR*D Report. Am J Psychiatry. 2006;13.

4. March JS, Silva S, Petrycki S, Curry J, Wells K, Fairbank J, et al. The Treatment for Adolescents With Depression Study (TADS): long-term effectiveness and safety outcomes. Arch Gen Psychiatry. 2007 Oct;64(10):1132–43. doi:10.1001/archpsyc.64.10.1132 PubMed PMID: 17909125.

5. Avenevoli S, Swendsen J, He JP, Burstein M, Merikangas KR. Major Depression in the National Comorbidity Survey–Adolescent Supplement: Prevalence, Correlates, and Treatment. J Am Acad Child Adolesc Psychiatry. 2015 Jan 1;54(1):37–44.e2. doi:10.1016/j.jaac.2014.10.010

6. Brenhouse HC, Schwarz JM. Immunoadolescence: Neuroimmune development and adolescent behavior. Neurosci Biobehav Rev. 2016 Nov 1;The Adolescent Brain70:288–99. doi:10.1016/j.neubiorev.2016.05.035

7. van Lang NDJ, Ferdinand RF, Verhulst FC. Predictors of future depression in early and late adolescence. J Affect Disord. 2007 Jan 1;97(1):137–44. doi:10.1016/j.jad.2006.06.007

8. Lewinsohn PM, Rohde P, Seeley JR, Klein DN, Gotlib IH. Psychosocial functioning of young adults who have experienced and recovered from major depressive disorder during adolescence. J Abnorm Psychol. 2003 Aug;112(3):353–63. doi:10.1037/0021-843X.112.3.353

9. McMakin DL, Olino TM, Porta G, Dietz LJ, Emslie G, Clarke G, et al. Anhedonia Predicts Poorer Recovery Among Youth With Selective Serotonin Reuptake Inhibitor Treatment– Resistant Depression. J Am Acad Child Adolesc Psychiatry. 2012 Apr 1;51(4):404–11. doi:10.1016/j.jaac.2012.01.011

10. Whitton AE, Kumar P, Treadway MT, Rutherford AV, Ironside ML, Foti D, et al. Distinct profiles of anhedonia and reward processing and their prospective associations with quality of life among individuals with mood disorders. Mol Psychiatry. 2023 Jul 4;1–10. doi:10.1038/s41380-023-02165-1

11. Pizzagalli DA, editor. Anhedonia: Preclinical, Translational, and Clinical Integration [Internet]. Vol. 58. Cham: Springer International Publishing; 2022 [cited 2025 Jun 23]. (Current Topics in Behavioral Neurosciences). Available from: https://link.springer.com/10.1007/978-3-031-09683-9 doi:10.1007/978-3-031-09683-9

12. Klein DF. Endogenomorphic depression. A conceptual and terminological revision. Arch Gen Psychiatry. 1974 Oct;31(4):447–54. doi:10.1001/archpsyc.1974.01760160005001 PubMed PMID: 4420562.

13. Dillon DG, Holmes AJ, Jahn AL, Bogdan R, Wald LL, Pizzagalli DA. Dissociation of neural regions associated with anticipatory versus consummatory phases of incentive processing. Psychophysiology. 2007 Sep 10;0(0):070915195953002-??? doi:10.1111/j.1469-8986.2007.00594.x

14. Treadway MT, Zald DH. Parsing Anhedonia: Translational Models of Reward-Processing Deficits in Psychopathology. Curr Dir Psychol Sci. 2013 Jun 1;22(3):244–9. doi:10.1177/0963721412474460 PubMed PMID: 24748727; PubMed Central PMCID: PMC3989147.

15. Mohebi A, Pettibone JR, Hamid AA, Wong JMT, Vinson LT, Patriarchi T, et al. Dissociable dopamine dynamics for learning and motivation. Nature. 2019 Jun;570(7759):7759. doi:10.1038/s41586-019-1235-y

16. Nguyen D, Naffziger EE, Berridge KC. Positive Affect: Nature and brain bases of liking and wanting. Curr Opin Behav Sci. 2021 Jun;39:72–8. doi:10.1016/j.cobeha.2021.02.013 PubMed PMID: 33748351; PubMed Central PMCID: PMC7978410.

17. Berridge KC, Kringelbach ML. Pleasure Systems in the Brain. Neuron. 2015;86(3). doi:10.1016/j.neuron.2015.02.018

18. Castro DC, Berridge KC. Opioid and orexin hedonic hotspots in rat orbitofrontal cortex and insula. Proc Natl Acad Sci U S A. 2017 Oct 24;114(43):E9125–34. doi:10.1073/pnas.1705753114 PubMed PMID: 29073109; PubMed Central PMCID: PMC5664503.

19. Kragel PA, Treadway MT, Admon R, Pizzagalli DA, Hahn EC. A mesocorticolimbic signature of pleasure in the human brain. Nat Hum Behav. 2023 Jun 29;1–12. doi:10.1038/s41562-023-01639-0

20. Felger JC, Treadway MT. Inflammation Effects on Motivation and Motor Activity: Role of Dopamine. Neuropsychopharmacol Off Publ Am Coll Neuropsychopharmacol. 2017;42(1):216–41. doi:10.1038/npp.2016.143 PubMed PMID: 27480574; PubMed Central PMCID: PMC5143486.

21. Vichaya EG, Dantzer R. Inflammation-induced motivational changes: perspective gained by evaluating positive and negative valence systems. Curr Opin Behav Sci. 2018 Aug 1;Apathy and Motivation22:90–5. doi:10.1016/j.cobeha.2018.01.008

22. Eisenberger NI, Moieni M, Inagaki TK, Muscatell KA, Irwin MR. In Sickness and in Health: The Co-Regulation of Inflammation and Social Behavior. Neuropsychopharmacol Off Publ Am Coll Neuropsychopharmacol. 2017;42(1):242–53. doi:10.1038/npp.2016.141 PubMed PMID: 27480575; PubMed Central PMCID: PMC5143485.

23. Nusslock R, Alloy LB, Brody GH, Miller GE. Annual Research Review: Neuroimmune network model of depression: a developmental perspective. J Child Psychol Psychiatry. 2024;65(4):538–67. doi:10.1111/jcpp.13961

24. Treadway MT, Cooper JA, Miller AH. Can’t or Won’t? Immunometabolic Constraints on Dopaminergic Drive. Trends Cogn Sci. 2019 May;23(5):435–48. doi:10.1016/j.tics.2019.03.003 PubMed PMID: 30948204.

25. Ben-Shaanan TL, Azulay-Debby H, Dubovik T, Starosvetsky E, Korin B, Schiller M, et al. Activation of the reward system boosts innate and adaptive immunity. Nat Med. 2016;22(8):940–4.

26. Ben-Shaanan TL, Schiller M, Azulay-Debby H, Korin B, Boshnak N, Koren T, et al. Modulation of anti-tumor immunity by the brain’s reward system. Nat Commun. 2018 Jul 13;9(1):1. doi:10.1038/s41467-018-05283-5

27. Mehta ND, Stevens JS, Li Z, Gillespie CF, Fani N, Michopoulos V, et al. Inflammation, reward circuitry and symptoms of anhedonia and PTSD in trauma-exposed women. Soc Cogn Affect Neurosci. 2020 Nov 10;15(10):1046–55. doi:10.1093/scan/nsz100

28. Bekhbat M, Li Z, Dunlop BW, Treadway MT, Mehta ND, Revill KP, et al. Sustained effects of repeated levodopa (L-DOPA) administration on reward circuitry, effort-based motivation, and anhedonia in depressed patients with higher inflammation. Brain Behav Immun. 2025 Mar 1;125:240–8. doi:10.1016/j.bbi.2024.12.026

29. Treadway MT, Etuk SM, Cooper JA, Hossein S, Hahn E, Betters SA, et al. A randomized proof-of-mechanism trial of TNF antagonism for motivational deficits and related corticostriatal circuitry in depressed patients with high inflammation. Mol Psychiatry. 2024;1–11.

30. Larsen B, Olafsson V, Calabro F, Laymon C, Tervo-Clemmens B, Campbell E, et al. Maturation of the human striatal dopamine system revealed by PET and quantitative MRI. Nat Commun. 2020 Feb 12;11(1):846. doi:10.1038/s41467-020-14693-3

31. Graham JM. Brain iron deposition in Parkinson’s disease imaged using the PRIME magnetic resonance sequence. Brain. 2000;123(12):2423–31.

32. Sedlacik J, Boelmans K, Löbel U, Holst B, Siemonsen S, Fiehler J. Reversible, irreversible and effective transverse relaxation rates in normal aging brain at 3T. NeuroImage. 2014;84:1032.

33. Peterson ET, Kwon D, Luna B, Larsen B, Prouty D, De Bellis MD, et al. Distribution of brain iron accrual in adolescence: Evidence from cross-sectional and longitudinal analysis. Hum Brain Mapp. 2019;40(5):1480–95.

34. Larsen B, Luna B. In vivo evidence of neurophysiological maturation of the human adolescent striatum. Dev Cogn Neurosci. 2015;12:74–85.

35. Sinclair D, Purves-Tyson TD, Allen KM, Weickert CS. Impacts of stress and sex hormones on dopamine neurotransmission in the adolescent brain. Psychopharmacology (Berl). 2014 Apr;231(8):1581–99. doi:10.1007/s00213-013-3415-z

36. Andersen SL, Teicher MH. Sex differences in dopamine receptors and their relevance to ADHD. Neurosci Biobehav Rev. 2000;24(1):137–41.

37. Reynolds LM, Flores C. Mesocorticolimbic Dopamine Pathways Across Adolescence: Diversity in Development. Front Neural Circuits [Internet]. 2021 [cited 2026 Apr 28];15. Available from: https://search.ebscohost.com/login.aspx?direct=trueCprofile=ehostCscope=siteCauthtype=crawlerCjrnl=16625110CAN=152375693Ch=3UBT8%2FoV3oCzIXDdfbNz0hwuHHB8KoL1OTAT1rinIfHiShvgSLFw8axuWgHxZkDwi0AXDAUljn5MsXn8r89nRQ%3D%3DCcrl=c

38. Snaith RP, Hamilton M, Morley S, Humayan A, Hargreaves D, Trigwell P. A Scale for the Assessment of Hedonic Tone the Snaith–Hamilton Pleasure Scale. Br J Psychiatry. 1995 Jul;167(1):99–103. doi:10.1192/bjp.167.1.99

39. Borsini A, Wallis ASJ, Zunszain P, Pariante CM, Kempton MJ. Characterizing anhedonia: A systematic review of neuroimaging across the subtypes of reward processing deficits in depression. Cogn Affect Behav Neurosci. 2020 Aug;20(4):816–41. doi:10.3758/s13415-020-00804-6

40. Cooper JA, Arulpragasam AR, Treadway MT. Anhedonia in depression: biological mechanisms and computational models. Curr Opin Behav Sci. 2018 Aug 1;Apathy and Motivation22:128–35. doi:10.1016/j.cobeha.2018.01.024

41. Khazanov GK, Ruscio AM, Forbes CN. The Positive Valence Systems Scale: Development and Validation. Assessment. 2020 Jul 1;27(5):1045–69. doi:10.1177/1073191119869836

42. First MB, Williams JB, Karg RS, Spitzer RL. Structured clinical interview for DSM-5— Research version (SCID-5 for DSM-5, research version; SCID-5-RV). Arlingt VA Am Psychiatr Assoc. 2015;(1–94).

43. Montgomery SA, Åsberg M. A New Depression Scale Designed to be Sensitive to Change. Br J Psychiatry. 1979 Apr;134(4):382–9. doi:10.1192/bjp.134.4.382

44. Haacke EM, Cheng NYC, House MJ, Liu Q, Neelavalli J, Ogg RJ, et al. Imaging iron stores in the brain using magnetic resonance imaging. Magn Reson Imaging. 2005 Jan;23(1):1–25. doi:10.1016/j.mri.2004.10.001 PubMed PMID: 15733784.

45. Costi S, Morris LS, Collins A, Fernandez NF, Patel M, Xie H, et al. Peripheral immune cell reactivity and neural response to reward in patients with depression and anhedonia. Transl Psychiatry. 2021;11(1):565.

46. Horn JL. A rationale and test for the number of factors in factor analysis. Psychometrika. 1965;30(2):179–85.

47. Ma Y. Neuropsychological mechanism underlying antidepressant effect: a systematic meta-analysis. Mol Psychiatry. 2015 Mar;20(3):311–9. doi:10.1038/mp.2014.24

48. O’Connor MF, Bower JE, Cho HJ, Creswell JD, Dimitrov S, Hamby ME, et al. To assess, to control, to exclude: Effects of biobehavioral factors on circulating inflammatory markers. Brain Behav Immun. 2009 Oct 1;23(7):887–97. doi:10.1016/j.bbi.2009.04.005

49. Stevens JS, Hamann S. Sex differences in brain activation to emotional stimuli: A meta-analysis of neuroimaging studies. Neuropsychologia. 2012 Jun 1;50(7):1578–93. doi:10.1016/j.neuropsychologia.2012.03.011

50. Leighton SP, Nerurkar L, Krishnadas R, Johnman C, Graham GJ, Cavanagh J. Chemokines in depression in health and in inflammatory illness: a systematic review and meta-analysis. Mol Psychiatry. 2018 Jan;23(1):48–58. doi:10.1038/mp.2017.205

51. Zhang E, Huang Z, Zang Z, Qiao X, Yan J, Shao X. Identifying circulating biomarkers for major depressive disorder. Front Psychiatry. 2023 Aug 4;14. doi:10.3389/fpsyt.2023.1230246

52. Toenders YJ, Laskaris L, Davey CG, Berk M, Milaneschi Y, Lamers F, et al. Inflammation and depression in young people: a systematic review and proposed inflammatory pathways. Mol Psychiatry. 2022 Jan;27(1):315–27. doi:10.1038/s41380-021-01306-8

53. Boyle CC, Bower JE, Eisenberger NI, Irwin MR. Stress to inflammation and anhedonia: Mechanistic insights from preclinical and clinical models. Neurosci Biobehav Rev. 2023 Sep;152:105307. doi:10.1016/j.neubiorev.2023.105307 PubMed PMID: 37419230.

54. Pizzagalli DA. Toward a Better Understanding of the Mechanisms and Pathophysiology of Anhedonia: Are We Ready for Translation? Am J Psychiatry. 2022 Jul 1;179(7):458–69. doi:10.1176/appi.ajp.20220423

55. Dantzer R, O’Connor JC, Freund GG, Johnson RW, Kelley KW. From inflammation to sickness and depression: when the immune system subjugates the brain. Nat Rev Neurosci. 2008 Jan;9(1):46–56. doi:10.1038/nrn2297 PubMed PMID: 18073775; PubMed Central PMCID: PMC2919277.

56. Freed RD, Mehra LM, Laor D, Patel M, Alonso CM, Kim-Schulze S, et al. Anhedonia as a clinical correlate of inflammation in adolescents across psychiatric conditions. World J Biol Psychiatry. 2019;20(9):712–22.

57. Rengasamy M, Marsland A, McClain L, Kovats T, Walko T, Pan L, et al. Longitudinal relationships of cytokines, depression and anhedonia in depressed adolescents. Brain Behav Immun. 2021;91:74–80.

58. Rizvi SJ, Pizzagalli DA, Sproule BA, Kennedy SH. Assessing anhedonia in depression: Potentials and pitfalls. Neurosci Biobehav Rev. 2016 Jun 1;65:21–35. doi:10.1016/j.neubiorev.2016.03.004

59. Baune BT, Sampson E, Louise J, Hori H, Schubert KO, Clark SR, et al. No evidence for clinical efficacy of adjunctive celecoxib with vortioxetine in the treatment of depression: A 6-week double-blind placebo controlled randomized trial. Eur Neuropsychopharmacol. 2021 Dec 1;53:34–46. doi:10.1016/j.euroneuro.2021.07.092

60. Hellmann-Regen J, Clemens V, Grözinger M, Kornhuber J, Reif A, Prvulovic D, et al. Effect of Minocycline on Depressive Symptoms in Patients With Treatment-Resistant Depression: A Randomized Clinical Trial. JAMA Netw Open. 2022 Sep 14;5(9):e2230367. doi:10.1001/jamanetworkopen.2022.30367

61. Jarkas DA, Villeneuve AH, Daneshmend AZ, Villeneuve PJ, McQuaid RJ. Sex differences in the inflammation-depression link: A systematic review and meta-analysis. Brain Behav Immun. 2024;121:257–68.

62. Lasselin J, Lekander M, Axelsson J, Karshikoff B. Sex differences in how inflammation affects behavior: what we can learn from experimental inflammatory models in humans. Front Neuroendocrinol. 2018;50:91–106.

63. Andersen SL, Rutstein M, Benzo JM, Hostetter JC, Teicher MH. Sex differences in dopamine receptor overproduction and elimination. Neuroreport. 1997;8(6):1495–7.

64. Jorgensen NA, Muscatell KA, McCormick EM, Prinstein MJ, Lindquist KA, Telzer EH. Neighborhood disadvantage, race/ethnicity and neural sensitivity to social threat and reward among adolescents. Soc Cogn Affect Neurosci. 2023;18(1):nsac053.

65. Wiley CR, De Wayne PW, Sigrist C, Brownlow BN, Markser A, Hong S, et al. Differences in inflammation among black and white individuals: A systematic review and meta-analysis. Brain Behav Immun. 2025;127:269–86.

