## supplementary material for "Basal Ganglia Dopamine Availability and Immune Activation Interact and Relate to Anhedonia Severity among Youth with Depression"

### Supplemental Results

#### Results based on Rotated Principal Component Solution

##### *Secondary Analyses*

**Interaction between Dopamine Availability in Subregions of Basal Ganglia and General Immune Activation (Principal Component 1)**. As noted in the main text, secondary analyses were conducted to examine basal ganglia subregions for any significant interaction effects observed at the whole-basal ganglia level (i.e., SHAPS overall anhedonia and PVSS anticipatory anhedonia).

Interaction effects between dopamine availability in the caudate, putamen, and pallidum and immune principal component 1 (PC1) scores were observed on overall anhedonia as measured by the SHAPS. For participants in the top 68.97%, 68.97%, and 74.14% of immune PC1 scores, respectively, lower dopamine availability in the caudate ($B$=-2.029, $SE$=.673, $p$=.004, $\Delta R^2$=.128), putamen ($B$=-1.273, $SE$=.447, $p$=.007, $\Delta R^2$=.120), and pallidum ($B$=-.682, $SE$=.326, $p$=.042, $\Delta R^2$=.069) were associated with greater SHAPS overall anhedonia. We did not detect an interaction effect with the nucleus accumbens (NAc; $B$=.001, $SE$=.356, $p$=.999, $\Delta R^2$<.001).

We also detected an interaction effect between dopamine availability in the putamen and immune PC1 scores on anticipatory anhedonia. Specifically, for participants in the top 56.90% of immune PC1 scores, respectively, lower dopamine availability in the putamen ($B$=.935, $SE$=.382, $p$=.018, $\Delta R^2$=.078) was associated with greater PVSS anticipatory anhedonia. Such interaction effects were not observed with other subregions (caudate: *B*=.719, *SE*=.603, *p*=.239, $\Delta R^2$=.019; NAc: *B*=.048, *SE*=.305, *p*=.876, $\Delta R^2$<.001; pallidum: *B*=.542, SE=.278, p=.057, , $\Delta R^2$=.053).

***Exploratory Analysese***

**Other Immune Principal Components (PC2–PC5)**. The interaction effect between dopamine availability in the basal ganglia and PC4 on SHAPS overall anhedonia was significant, *B*=-.767, *SE*=.359, *p*=.038, $\Delta R^2$=.067, such that for participants in the top 60.34% of PC4 score, lower dopamine availability in the basal ganglia was associated with greater SHAPS overall anhedonia. Follow-up analyses probing the effect within subregions of basal ganglia observed an effect with putamen (*B*=-.764, *SE*=.288, *p*=.011, $\Delta R^2$=.100) and pallidum (*B*=-.472, *SE*=.207, *p*=.027, $\Delta R^2$=.076), such that for participants in the top 60.34% and 58.62% of immune PC4 scores, lower dopamine availability in the putamen and pallidum were associated with higher SHAPS overall anhedonia. Such interaction effects were not observed within other subregions (caudate: *B*=-.410, *SE*=.615, *p*=.508, $\Delta R^2$=.007; NAc: *B*=.387, *SE*=.350, *p*=.276, $\Delta R^2$=.020).

The interaction effect between dopamine availability in the basal ganglia and PC4 on anticipatory anhedonia was significant, *B*=.708, *SE*=.306, *p*=.025, $\Delta R^2$=.069, such that for participants in the top 37.93% of PC4 score, lower dopamine availability in the basal ganglia was associated with greater anticipatory anhedonia. Follow-up analyses probing the effect within subregions of basal ganglia observed an effect with putamen (*B*=.604, *SE*=.251, *p*=.020, $\Delta R^2$=.076) and pallidum (*B*=.509, *SE*=.174 *p*=.005, $\Delta R^2$=.107), such that for participants in the top 39.66% and 44.83% of immune PC4 scores, lower dopamine availability in the putamen and pallidum were associated with higher anticipatory anhedonia. Such interaction effects were not observed within other subregions (caudate: *B*=.573, *SE*=.520, *p*=.276, $\Delta R^2$=.017; NAc: *B*=-.123, *SE*=.323, *p*=.707, $\Delta R^2$=.002).

We did not detect interaction effects between dopamine availability in the basal ganglia and immune PC4 scores on PVSS overall anhedonia and consummatory anhedonia (*p's* > .05).

For immune PC2, PC3, and PC5 scores, there were no main effects or interaction effects for dopamine availability in the basal ganglia and immune PC scores on overall scores of anhedonia, anticipatory anhedonia, or consummatory anhedonia (*p's* > .05).

#### *Sensitivity Analyses*

Attention-deficit/hyperactivity disorder status was included as an additional covariate in primary analyses, and results remain unchanged (SHAPS overall anhedonia: B=-1.702, SE =.526, p =.002, $\Delta R^2$=.148; anticipatory anhedonia: B=1.124, SE=.461, p=.019, $\Delta R^2$=.078; PVSS overall anhedonia: B=.206, SE=.128, p=.114, $\Delta R^2$=.042; consummatory anhedonia: B=.760, SE=.577, p=.195, $\Delta R^2$=.032).

Sensitivity analyses using a complete-case approach (i.e., excluding outlier value rather than imputing them) yielded interaction effects between basal ganglia dopamine availability and Immune PC1 that were consistent in direction with the primary analyses, although statistical significance was attenuated (SHAPS overall anhedonia: B=.064, SE =.035, p =.084, $\Delta R^2$=.087; anticipatory anhedonia: B=.063, SE=.033, p=.072, $\Delta R^2$=.089; PVSS overall anhedonia: B=.001, SE=.008, p=.896, $\Delta R^2$<.001; consummatory anhedonia: B=.013, SE=.035, p=.726, $\Delta R^2$=.004), likely due to reduced sample size (N=35).

### Results based on Unrotated Principal Component Solution

Analyses were repeated using the unrotated principal component solution to assess whether results depended on the choice of rotation. Findings were consistent in direction with the primary analyses, indicating that results were not driven by the rotation method. Full statistical results are reported below.

#### *Primary Analyses*

**General Immune Activation (PC1) and Anhedonia**. Main effects were found for higher immune activation (PC1 score, characterized by highest loadings on inflammatory cytokines, chemokines, and tissue remodeling markers [e.g., LAP TGF-β1, TNFSF14, CD40, VEGFA, IL-18)) on more severe SHAPS overall anhedonia ($B$=3.483, $SE$=1.310, $p$=.011, $\Delta R^2$=.105) and PVSS anticipatory anhedonia ($B$=-2.859, $SE$=1.104, $p$=.013, $\Delta R^2$=.085). Immune activation was unrelated to PVSS overall anhedonia ($B$=-.004, $SE$=.047, $p$=.931, $\Delta R^2$<.001) and PVSS consummatory anhedonia ($B$=.067, $SE$=.209, $p$=.751, $\Delta R^2$=.002).

**Dopamine Availability in Basal Ganglia and Anhedonia**. We did not detect main effects for dopamine availability within the basal ganglia on SHAPS overall anhedonia ($B$=-.387, SE=.341, $p$=.262, $\Delta R^2$=.019), PVSS overall anhedonia ($B$=.111, $SE$=.082, $p$=.179, $\Delta R^2$=.031) or PVSS consummatory anhedonia ($B$=.378, $SE$=.364, $p$=.305, $\Delta R^2$=.020), except for PVSS anticipatory anhedonia ($B$=.623, $SE$=.287, $p$=.035, $\Delta R^2$=.060).

**Interaction of Dopamine Availability in Basal Ganglia and Immune Activation in Relation to Anhedonia**. There was an interaction effect between dopamine availability in the basal ganglia and immune activation on SHAPS overall anhedonia ($B$=-.299, $SE$=.109, $p$=.009, $\Delta R^2$=.111), such that for participants in the top 68.97% of levels of immune activation, lower dopamine availability was associated with more severe SHAPS overall anhedonia (Figure 2A). We did not detect an interaction effect for PVSS overall anhedonia ($B$=.042, $SE$=.026, $p$=.113, $\Delta R^2$=.042; Figure 2B).

There was also an interaction effect for PVSS anticipatory anhedonia (B=.235, $SE$=.092, $p$=.014, $\Delta R^2$=.084), such that for participants in the top 58.62% of levels of immune activation, lower dopamine availability was significantly associated with more severe PVSS anticipatory anhedonia (Figure 3A). We did not detect an interaction effect for PVSS consummatory anhedonia ($B$=.144, $SE$=.116, $p$=.220, $\Delta R^2$=.028; Figure 3B).

***Secondary Analyses***

**Interaction between Dopamine Availability in Subregions of Basal Ganglia and General Immune Activation (Principal Component 1)**. Secondary analyses were conducted to examine basal ganglia subregions for any interaction effects observed at the whole-basal ganglia level. Interaction effects between dopamine availability in the caudate, putamen, and pallidum and immune principal component 1 (PC1) scores were observed on overall anhedonia as measured by the SHAPS. For participants in the top 68.97%, 70.69%, and 74.14% of immune PC1 scores, respectively, lower dopamine availability in the caudate ($B$=-.341, $SE$=.159, $p$=.037, $\Delta R^2$=.071), putamen ($B$=-.242, $SE$=.090, $p$=.010, $\Delta R^2$=.109), and pallidum ($B$=-.129, $SE$=.061, $p$=.041, $\Delta R^2$=.070) were associated with greater SHAPS overall anhedonia. We did not detect an interaction effect with the nucleus accumbens (NAc; $B$=.079, $SE$=.087, $p$=.371, $\Delta R^2$=.016).

We also detected an interaction effect between dopamine availability in the putamen and pallidum and immune PC1 scores on anticipatory anhedonia. Specifically, for participants in the top 58.62% and 67.24% of immune PC1 scores, respectively, lower dopamine availability in the putamen (*B*=.187, *SE*=.076, *p*=.017, $\Delta R^2$=.079) and pallidum (*B*=.128, *SE*=.051, *p*=.016, $\Delta R^2$=.083) were associated with greater PVSS anticipatory anhedonia. Such interaction effects were not observed with other subregions (caudate: *B*=.155, *SE*=.136, *p*=.262, $\Delta R^2$=.018; NAc: *B*=.042, *SE*=.075, *p*=.583, $\Delta R^2$=.005).

***Exploratory Analysese***

**Other Immune Principal Components (PC2–PC5)**. Lower PC2 (characterized by higher positive loadings on 4E-BP1, AXIN1, SIRT2, STAMBP, and CASP-8, and higher negative loadings on TNF-β and CX3CL1) scores were associated with greater PVSS overall anhedonia (*B*=.169, *SE*=.068, *p*=.016, $\Delta R^2$=.092). The interaction effect between dopamine availability in the basal ganglia and PC2 on PVSS overall anhedonia was not significant, *B*=-.004, *SE*=.033, *p*=.911, $\Delta R^2$=.001. There were no main effect for PC2 score (*B*=-.408, *SE*=.316, *p*=.203, $\Delta R^2$= .027) or interaction effect between basal ganglia dopamine availability and PC2 on SHAPS overall anhedonia (*B*=-.102, *SE*=.155, *p*=.513, $\Delta R^2$=.007). The main effect of PC2 score on higher PVSS anticipatory anhedonia was not significant, *B*= .413, *SE*=.261, *p*=.121, $\Delta R^2$=.034. The interaction effect between basal ganglia dopamine availability and PC2 on PVSS anticipatory anhedonia also was not significant, *B*=.128, *SE*=.128, *p*=.320, $\Delta R^2$=.014). Lower PC2 scores were associated with higher PVSS consummatory anhedonia, *B*=.739, *SE*=.303, *p*=.019, $\Delta R^2$=.098. The interaction effect between basal ganglia dopamine availability and PC2 on consummatory anhedonia were not significant, *B*=-.047, *SE*=.149, *p*=.756, $\Delta R^2$=.002).

For immune PC3 (higher positive loadings on TWEAK, and EN-RAGE, and negative loadings on ST1A1, SIRT2, AXIN1) and PC4 (higher positive loadings on TRAIL, uPA, CD8A, and TNF, and negative loadings on ST1A1) scores, there were no main effects or interaction effects for dopamine availability in basal ganglia and immune PC scores on overall scores of anhedonia, anticipatory anhedonia, or consummatory anhedonia (*p's* > .05).

There were no main effects of PC5 (characterized by higher positive loadings on DNER, CX3CL1, TNF-β, and ADA, and higher negative loadings on CD8A, IL-6, OSM, and CCL23) score on overall anhedonia, anticipatory anhedonia, or consummatory anhedonia (*p's* > .05). An interaction effect between basal ganglia dopamine availability and immune PC5 scores was detected for SHAPS overall anhedonia, $B$=.564, $SE$=.275, $p$=.046, $\Delta R^2$=.064, such that for participants in the bottom 32.76% of PC5 score, lower dopamine availability in the basal ganglia was associated with greater SHAPS overall anhedonia. Follow-up analyses probing the effect within subregions of basal ganglia observed an effect with putamen, $B$=.488, $SE$=.232, $p$=.041, $\Delta R^2$=.067, such that for participants in the bottom 27.59% of immune PC5 scores, lower dopamine availability in the putamen was associated with higher SHAPS overall anhedonia. Such interaction effects were not observed within other subregions (caudate: $B$=.527, $SE$=.375, $p$=.167, $\Delta R^2$=.031; pallidum: $B$=.246, $SE$=.162, $p$=.135, $\Delta R^2$=.037; NAc: $B$=-.043, $SE$=.202, $p$=.832, $\Delta R^2$=.001). We did not detect interaction effects between dopamine availability in the basal ganglia and immune PC5 scores on PVSS overall anhedonia, anticipitary anhedonia, and consummatory anhedonia (*p's* > .05).

**Supplementary Table 1.** Summary of NPX values for each of the immune protein included in PCA.

| Immune Proteins | Mean NPX units, log2 scale | Standard Deviation |
|---|---|---|
| IL8 | 5.91 | 0.54 |
| VEGFA | 10.82 | 0.42 |
| CD8A | 6.69 | 1.29 |
| CDCP1 | 2.46 | 0.44 |
| CD244 | 6.59 | 0.36 |
| IL7 | 1.34 | 0.56 |
| OPG | 10.12 | 0.33 |
| LAP TGF-beta-1 | 6.53 | 0.40 |
| uPA | 10.30 | 0.39 |
| IL6 | 2.71 | 0.89 |
| IL-17C | 1.96 | 0.87 |
| MCP-1 | 10.50 | 0.34 |
| CXCL11 | 8.45 | 0.86 |
| AXIN1 | 2.93 | 1.24 |
| TRAIL | 7.41 | 0.37 |
| CXCL9 | 6.70 | 0.73 |
| CST5 | 6.87 | 0.53 |
| OSM | 4.11 | 1.03 |
| CXCL1 | 10.33 | 0.76 |
| CCL4 | 6.01 | 0.54 |
| CD6 | 5.79 | 0.58 |
| SCF | 9.22 | 0.38 |
| IL18 | 8.88 | 0.56 |
| SLAMF1 | 2.36 | 0.38 |
| TGF-alpha | 1.49 | 0.51 |
| MCP-4 | 14.68 | 0.70 |
| CCL11 | 6.50 | 0.68 |
| TNFSF14 | 3.86 | 0.71 |
| FGF-23 | 1.93 | 0.69 |
| MMP-1 | 11.54 | 1.13 |
| LIF-R | 4.30 | 0.29 |
| FGF-21 | 3.40 | 1.49 |
| CCL19 | 11.27 | 1.04 |
| IL-10RB | 6.23 | 0.30 |
| IL-18R1 | 8.48 | 0.41 |
| PD-L1 | 6.57 | 0.32 |
| CXCL5 | 11.97 | 1.20 |
| TRANCE | 5.53 | 0.65 |
| HGF | 8.84 | 0.51 |
| IL-12B | 6.56 | 0.61 |
| MMP-10 | 9.10 | 0.70 |
| IL10 | 3.72 | 0.63 |
| TNF | 3.25 | 0.58 |
| CCL23 | 11.81 | 0.39 |
| CD5 | 5.29 | 0.32 |
| CCL3 | 5.49 | 0.64 |
| Flt3L | 8.85 | 0.46 |
| CXCL6 | 8.59 | 0.70 |
| CXCL10 | 7.99 | 0.71 |
| 4E-BP1 | 4.30 | 2.35 |

| Immune Proteins | Mean NPX units, log2 scale | Standard Deviation |
|---|---|---|
| SIRT2 | 4.90 | 1.42 |
| CCL28 | 2.65 | 0.83 |
| DNER | 8.68 | 0.24 |
| EN-RAGE | 2.80 | 1.05 |
| CD40 | 11.35 | 0.42 |
| IFN-gamma | 5.76 | 0.78 |
| FGF-19 | 7.59 | 0.89 |
| MCP-2 | 8.84 | 0.65 |
| CASP-8 | 4.00 | 0.86 |
| CCL25 | 5.66 | 0.58 |
| CX3CL1 | 4.05 | 0.70 |
| TNFRSF9 | 5.35 | 0.42 |
| NT-3 | 1.94 | 0.42 |
| TWEAK | 9.33 | 0.37 |
| CCL20 | 6.39 | 0.93 |
| ST1A1 | 7.01 | 1.07 |
| STAMBP | 6.02 | 0.89 |
| ADA | 5.60 | 0.38 |
| TNFB | 3.98 | 0.96 |
| CSF-1 | 10.29 | 0.21 |

**Note.** See Supplementary List of Abbreviations for full protein names.

**Supplementary Table 2.** Number and percentage of samples imputed for each immune marker.

| | Markers | samples imputed (n) | samples imputed (%) |
|---|---|---|---|
| 1 | MCP-1 | 3 | 3.8 |
| 2 | VEGFA | 3 | 3.8 |
| 3 | CCL19 | 2 | 2.53 |
| 4 | CCL3 | 2 | 2.53 |
| 5 | CD6 | 2 | 2.53 |
| 6 | EN-RAGE | 2 | 2.53 |
| 7 | IFN-gamma | 2 | 2.53 |
| 8 | IL-12B | 2 | 2.53 |
| 9 | IL7 | 2 | 2.53 |
| 10 | IL8 | 2 | 2.53 |
| 11 | MCP-2 | 2 | 2.53 |
| 12 | NT-3 | 2 | 2.53 |
| 13 | OPG | 2 | 2.53 |
| 14 | TGF-alpha | 2 | 2.53 |
| 15 | ADA | 1 | 1.27 |
| 16 | CCL20 | 1 | 1.27 |
| 17 | CCL23 | 1 | 1.27 |
| 18 | CCL28 | 1 | 1.27 |
| 19 | CCL4 | 1 | 1.27 |
| 20 | CD5 | 1 | 1.27 |
| 21 | CDCP1 | 1 | 1.27 |
| 22 | CST5 | 1 | 1.27 |
| 23 | CXCL9 | 1 | 1.27 |
| 24 | DNER | 1 | 1.27 |
| 25 | Flt3L | 1 | 1.27 |
| 26 | HGF | 1 | 1.27 |
| 27 | IL-17C | 1 | 1.27 |
| 28 | IL10 | 1 | 1.27 |
| 29 | IL18 | 1 | 1.27 |
| 30 | IL6 | 1 | 1.27 |
| 31 | MCP-4 | 1 | 1.27 |
| 32 | MMP-10 | 1 | 1.27 |
| 33 | PD-L1 | 1 | 1.27 |
| 34 | SLAMF1 | 1 | 1.27 |
| 35 | ST1A1 | 1 | 1.27 |
| 36 | TNF | 1 | 1.27 |
| 37 | TRANCE | 1 | 1.27 |
| 38 | TWEAK | 1 | 1.27 |

**Note**. Values indicate the number and percentage of samples with imputed protein expression values. Markers not shown (n=32) had no imputed values.

**Supplementary Table 3.** Oblimin-rotated loadings of immune proteins across the five retained principal components.

| Immune Proteins | Principal Component 1 | Principal Component 2 | Principal Component 3 | Principal Component 4 | Principal Component 5 |
|---|---|---|---|---|---|
| HGF | 0.874 | -0.189 | 0.09 | -0.008 | -0.131 |
| IL7 | 0.818 | 0.285 | -0.13 | -0.07 | 0.153 |
| VEGFA | 0.809 | 0.08 | 0.104 | 0.034 | -0.112 |
| TGF-alpha | 0.807 | -0.27 | 0.085 | 0.095 | 0.038 |
| TNFSF14 | 0.806 | 0.173 | 0.121 | 0.018 | -0.101 |
| OSM | 0.789 | -0.088 | -0.009 | 0.057 | -0.325 |
| MMP-1 | 0.694 | 0.177 | -0.098 | 0.06 | 0.195 |
| IL8 | 0.691 | 0.139 | 0.093 | -0.127 | 0.148 |
| EN-RAGE | 0.689 | -0.238 | -0.103 | 0.065 | 0.052 |
| MCP-4 | 0.589 | 0.304 | -0.016 | -0.07 | 0.301 |
| CCL28 | 0.584 | 0.121 | -0.071 | -0.006 | 0.382 |
| LAP TGF-beta-1 | 0.578 | 0.171 | 0.154 | 0.335 | 0.17 |
| MCP-1 | 0.537 | -0.145 | 0.246 | 0.237 | 0.288 |
| IL6 | 0.51 | 0.115 | 0.197 | -0.095 | -0.569 |
| MCP-2 | 0.507 | 0.075 | 0.133 | -0.032 | 0.259 |
| CXCL5 | 0.479 | 0.562 | -0.216 | -0.068 | 0.34 |
| CSF-1 | 0.444 | 0.009 | 0.514 | -0.016 | -0.316 |
| IL-10RB | 0.411 | -0.097 | 0.396 | -0.028 | -0.287 |
| IL-18R1 | 0.411 | 0.019 | 0.287 | -0.12 | -0.146 |
| CD40 | 0.41 | 0.504 | 0.294 | 0.065 | -0.075 |
| CXCL6 | 0.402 | 0.56 | 0.053 | -0.01 | 0.267 |
| CCL4 | 0.369 | 0.096 | 0.44 | -0.048 | 0.224 |
| TWEAK | 0.356 | -0.32 | 0.156 | 0.031 | 0.568 |
| OPG | 0.345 | -0.038 | 0.089 | -0.01 | -0.283 |
| PD-L1 | 0.294 | 0.109 | 0.565 | -0.05 | 0.024 |
| CCL25 | 0.284 | -0.001 | -0.017 | 0.213 | 0.414 |
| CXCL1 | 0.248 | 0.63 | -0.148 | 0.059 | 0.199 |
| CASP-8 | 0.243 | 0.678 | 0.073 | 0.349 | -0.015 |
| IL18 | 0.239 | 0.186 | 0.6 | 0.049 | -0.007 |
| CCL11 | 0.238 | -0.062 | 0.06 | 0.576 | 0.453 |
| LIF-R | 0.234 | -0.472 | 0.284 | -0.039 | 0.405 |
| CXCL11 | 0.214 | 0.621 | 0.207 | -0.231 | 0.134 |
| CCL3 | 0.211 | 0.023 | 0.685 | 0.029 | -0.093 |
| CD8A | 0.196 | -0.151 | 0.102 | 0.845 | -0.103 |
| CCL23 | 0.182 | -0.281 | 0.339 | 0.133 | -0.39 |
| TRANCE | 0.171 | -0.004 | 0.053 | 0.067 | -0.16 |
| CX3CL1 | 0.166 | -0.333 | 0.061 | -0.707 | 0.272 |
| CDCP1 | 0.109 | 0.097 | 0.667 | -0.023 | -0.02 |
| MMP-10 | 0.085 | -0.16 | 0.217 | -0.201 | 0.027 |
| uPA | 0.082 | -0.17 | 0.197 | 0.642 | 0.263 |
| IL-12B | 0.065 | -0.042 | 0.629 | -0.06 | -0.309 |
| ADA | 0.064 | 0.343 | 0.289 | -0.041 | 0.392 |
| TNFRSF9 | 0.054 | -0.049 | 0.699 | 0.173 | -0.156 |
| TNFB | 0.038 | -0.057 | 0.305 | -0.897 | -0.011 |
| IL-17C | 0.036 | 0.142 | 0.193 | 0.512 | 0.144 |
| STAMBP | 0.035 | 0.922 | 0.054 | 0.082 | -0.059 |
| SLAMF1 | 0.031 | 0.133 | 0.578 | 0.074 | 0.18 |
| FGF-21 | 0 | 0.198 | 0.152 | 0.233 | -0.051 |

| | | | | | |
|---|---|---|---|---|---|
| CD5 | -0.016 | -0.102 | 0.787 | 0.147 | -0.031 |
| CD6 | -0.028 | -0.097 | 0.73 | -0.023 | 0.251 |
| DNER | -0.035 | -0.102 | 0.049 | 0.021 | 0.686 |
| CD244 | -0.037 | 0.24 | 0.738 | -0.106 | 0.177 |
| TNF | -0.042 | 0.059 | 0.598 | 0.453 | -0.009 |
| CCL20 | -0.043 | 0.163 | 0.388 | -0.336 | 0.119 |
| SIRT2 | -0.044 | 0.904 | 0.066 | 0.156 | -0.091 |
| CST5 | -0.05 | -0.018 | 0.195 | 0.106 | 0.389 |
| CXCL9 | -0.054 | 0.026 | 0.676 | -0.183 | 0.034 |
| TRAIL | -0.06 | 0 | 0.317 | 0.623 | 0.264 |
| 4E-BP1 | -0.067 | 0.326 | 0.009 | 0.849 | 0.037 |
| IFN-gamma | -0.074 | -0.016 | 0.685 | -0.013 | -0.032 |
| CXCL10 | -0.078 | 0.002 | 0.708 | -0.333 | 0.028 |
| NT-3 | -0.085 | 0.15 | 0.052 | 0.338 | 0.373 |
| AXIN1 | -0.086 | 0.878 | 0.066 | 0.238 | -0.069 |
| Flt3L | -0.098 | 0.027 | 0.273 | 0.07 | 0.192 |
| SCF | -0.102 | -0.226 | -0.053 | 0.219 | 0.527 |
| ST1A1 | -0.137 | 0.792 | 0.065 | -0.344 | -0.111 |
| IL10 | -0.149 | 0.193 | 0.649 | 0.125 | -0.01 |
| CCL19 | -0.164 | -0.076 | 0.353 | 0.12 | 0.067 |
| FGF-23 | -0.224 | 0.356 | 0.291 | 0.211 | -0.002 |
| FGF-19 | -0.27 | 0.063 | 0.38 | 0.019 | 0.371 |

**Note.** Proteins are listed in descending order of their loadings on Principal Component 1 (from highest positive to most negative). Loadings are color-coded by direction and magnitude using a continuous gradient centered at 0 (blue = negative, red = positive, white ≈ 0), with values ≥ |0.5| displayed at maximum color intensity. See Supplementary List of Abbreviations for full protein names.

**Supplementary Table 4.** Unrotated loadings of immune proteins across the retained principal components.

| Immune Proteins | Principal Component 1 | Principal Component 2 | Principal Component 3 | Principal Component 4 | Principal Component 5 |
|---|---|---|---|---|---|
| LAP TGF-beta-1 | 0.193 | 0.069 | 0.1 | 0.01 | -0.05 |
| TNFSF14 | 0.189 | -0.032 | 0.069 | -0.141 | -0.108 |
| CD40 | 0.184 | 0.057 | -0.089 | -0.096 | -0.031 |
| VEGFA | 0.177 | -0.049 | 0.088 | -0.125 | -0.124 |
| IL18 | 0.173 | -0.059 | -0.096 | 0.034 | 0.009 |
| IL7 | 0.169 | 0.038 | 0.145 | -0.195 | 0.008 |
| CCL4 | 0.168 | -0.066 | 0.016 | 0.011 | 0.093 |
| MCP-1 | 0.167 | -0.031 | 0.16 | 0.072 | 0.003 |
| PD-L1 | 0.164 | -0.094 | -0.064 | 0.011 | 0.03 |
| CXCL6 | 0.163 | 0.125 | 0.011 | -0.123 | 0.114 |
| IL8 | 0.163 | -0.038 | 0.105 | -0.133 | 0.033 |
| CCL3 | 0.159 | -0.119 | -0.099 | 0.066 | -0.026 |
| CASP-8 | 0.158 | 0.213 | -0.086 | -0.053 | -0.055 |
| MCP-4 | 0.156 | 0.052 | 0.112 | -0.131 | 0.098 |
| HGF | 0.155 | -0.128 | 0.147 | -0.104 | -0.152 |
| CSF-1 | 0.154 | -0.141 | -0.082 | -0.03 | -0.139 |
| MMP-1 | 0.152 | 0.048 | 0.153 | -0.115 | -0.001 |
| TGF-alpha | 0.151 | -0.105 | 0.198 | -0.037 | -0.109 |
| CD244 | 0.149 | -0.068 | -0.149 | 0.066 | 0.159 |
| CDCP1 | 0.144 | -0.1 | -0.121 | 0.059 | 0.03 |
| MCP-2 | 0.142 | -0.018 | 0.109 | -0.05 | 0.07 |
| SLAMF1 | 0.137 | -0.033 | -0.076 | 0.098 | 0.093 |
| TNF | 0.137 | 0.016 | -0.099 | 0.203 | -0.069 |
| CCL28 | 0.136 | 0.039 | 0.178 | -0.083 | 0.097 |
| CXCL11 | 0.134 | 0.066 | -0.104 | -0.154 | 0.147 |
| CD5 | 0.132 | -0.125 | -0.116 | 0.172 | -0.011 |
| CXCL5 | 0.131 | 0.159 | 0.085 | -0.184 | 0.134 |
| TNFRSF9 | 0.13 | -0.107 | -0.127 | 0.134 | -0.074 |
| CD6 | 0.125 | -0.125 | -0.055 | 0.151 | 0.139 |
| OSM | 0.125 | -0.085 | 0.092 | -0.127 | -0.232 |
| CCL11 | 0.123 | 0.126 | 0.178 | 0.181 | 0.016 |
| ADA | 0.121 | 0.058 | -0.022 | 0.006 | 0.201 |
| STAMBP | 0.117 | 0.227 | -0.192 | -0.14 | 0.032 |
| IL-10RB | 0.117 | -0.146 | -0.041 | -0.025 | -0.136 |
| IL10 | 0.114 | -0.027 | -0.172 | 0.121 | 0.033 |
| IL-18R1 | 0.112 | -0.107 | -0.019 | -0.069 | -0.055 |
| TRAIL | 0.109 | 0.11 | 0.028 | 0.247 | -0.02 |
| CXCL1 | 0.107 | 0.194 | -0.006 | -0.132 | 0.082 |
| SIRT2 | 0.106 | 0.238 | -0.208 | -0.107 | 0.009 |
| IL-17C | 0.104 | 0.125 | 0.007 | 0.148 | -0.046 |
| AXIN1 | 0.103 | 0.254 | -0.201 | -0.073 | 0.001 |

| | | | | | |
|---|---|---|---|---|---|
| IL6 | 0.101 | -0.107 | -0.097 | -0.153 | -0.235 |
| TWEAK | 0.1 | -0.069 | 0.228 | 0.091 | 0.162 |
| IFN-gamma | 0.099 | -0.118 | -0.138 | 0.111 | 0.034 |
| CD8A | 0.099 | 0.108 | 0.073 | 0.22 | -0.267 |
| CXCL9 | 0.098 | -0.139 | -0.139 | 0.064 | 0.101 |
| uPA | 0.098 | 0.084 | 0.109 | 0.241 | -0.06 |
| EN-RAGE | 0.097 | -0.064 | 0.203 | -0.058 | -0.091 |
| CCL25 | 0.094 | 0.069 | 0.159 | 0.053 | 0.082 |
| IL-12B | 0.092 | -0.16 | -0.161 | 0.045 | -0.084 |
| 4E-BP1 | 0.092 | 0.27 | -0.02 | 0.181 | -0.152 |
| CXCL10 | 0.084 | -0.182 | -0.151 | 0.035 | 0.136 |
| LIF-R | 0.066 | -0.149 | 0.173 | 0.12 | 0.122 |
| FGF-23 | 0.062 | 0.098 | -0.149 | 0.072 | 0.017 |
| FGF-21 | 0.062 | 0.07 | -0.063 | 0.044 | -0.051 |
| OPG | 0.059 | -0.073 | -0.011 | -0.065 | -0.144 |
| CST5 | 0.055 | 0.024 | 0.052 | 0.109 | 0.143 |
| CCL20 | 0.054 | -0.084 | -0.098 | -0.037 | 0.165 |
| NT-3 | 0.054 | 0.142 | 0.05 | 0.126 | 0.091 |
| CCL23 | 0.05 | -0.144 | -0.05 | 0.065 | -0.209 |
| Flt3L | 0.049 | 0.001 | -0.024 | 0.089 | 0.086 |
| FGF-19 | 0.045 | 0.006 | -0.043 | 0.132 | 0.196 |
| TRANCE | 0.039 | -0.02 | -0.009 | -0.019 | -0.094 |
| ST1A1 | 0.037 | 0.106 | -0.235 | -0.208 | 0.12 |
| DNER | 0.036 | 0.036 | 0.158 | 0.108 | 0.263 |
| CCL19 | 0.034 | -0.033 | -0.053 | 0.128 | 0.028 |
| MMP-10 | 0.024 | -0.12 | 0.003 | -0.005 | 0.048 |
| SCF | -0.003 | 0.055 | 0.164 | 0.159 | 0.145 |
| TNFB | -0.016 | -0.263 | -0.089 | -0.185 | 0.216 |
| CX3CL1 | -0.031 | -0.23 | 0.104 | -0.115 | 0.233 |

***Note.*** Proteins are listed in descending order of their loadings on Principal Component 1 (from highest positive to most negative). Loadings are color-coded by direction and magnitude using a continuous gradient centered at 0 (blue = negative, red = positive, white ≈ 0), with values ≥ |0.5| displayed at maximum color intensity. See Supplementary List of Abbreviations for full protein names.

### List of Abbreviations

Abbreviations for immune proteins included in Supplementary Tables 3–4 are listed below (in alphabetical order).

4E-BP1, eukaryotic translation initiation factor 4E–binding protein 1;
ADA, adenosine deaminase;
AXIN1, axis inhibition protein 1;
CASP-8, caspase-8;
CCL, C-C motif chemokine ligand;
CD, cluster of differentiation;
CD5, cluster of differentiation 5;
CD6, cluster of differentiation 6;
CD8A, cluster of differentiation 8 alpha chain;
CD40, cluster of differentiation 40;
CD244, cluster of differentiation 244;
CDCP1, CUB domain–containing protein 1;
CSF-1, colony-stimulating factor 1;
CST5, cystatin D;
CX3CL1, C-X3-C motif chemokine ligand 1 (fractalkine);
CXCL, C-X-C motif chemokine ligand;
DNER, delta/notch-like EGF-related receptor;
EN-RAGE, extracellular newly identified receptor for advanced glycation end products binding protein (S100A12);
FGF, fibroblast growth factor;
FGF-19, fibroblast growth factor 19;
FGF-21, fibroblast growth factor 21;
FGF-23, fibroblast growth factor 23;
Flt3L, Fms-like tyrosine kinase 3 ligand;
HGF, hepatocyte growth factor;
IFN-γ, interferon gamma;
IL, interleukin;
LAP TGF-β-1, latency-associated peptide transforming growth factor beta-1;
LIF-R, leukemia inhibitory factor receptor;
MCP, monocyte chemoattractant protein;
MMP, matrix metalloproteinase;
MMP-1, matrix metalloproteinase-1;
MMP-10, matrix metalloproteinase-10;
NT-3, neurotrophin-3;
OPG, osteoprotegerin;
OSM, oncostatin M;
PD-L1, programmed death-ligand 1;
SCF, stem cell factor;
SIRT2, sirtuin 2;

SLAMF1, signaling lymphocytic activation molecule family member 1;
ST1A1, sulfotransferase family 1A member 1;
STAMBP, signal transducing adaptor molecule binding protein;
TGF-α, transforming growth factor alpha;
TNF, tumor necrosis factor;
TNFB, tumor necrosis factor beta;
TNFRSF9, tumor necrosis factor receptor superfamily member 9;
TNFSF14, tumor necrosis factor superfamily member 14;
TRAIL, TNF-related apoptosis-inducing ligand;
TRANCE, TNF-related activation-induced cytokine (RANKL);
TWEAK, TNF-like weak inducer of apoptosis;
uPA, urokinase-type plasminogen activator;
VEGFA, vascular endothelial growth factor A.
